# Active integrins regulate white adipose tissue insulin sensitivity and brown fat thermogenesis

**DOI:** 10.1101/2020.06.29.178020

**Authors:** Francisco Javier Ruiz-Ojeda, Jiefu Wang, Theresa Bäcker, Martin Krueger, Samira Zamani, Simon Rosowski, Tim Gruber, Annette Feuchtinger, Tim J. Schulz, Reinhard Fässler, Timo D. Müller, Cristina García-Cáceres, Matthias Meier, Matthias Blüher, Siegfried Ussar

## Abstract

Reorganization of the extracellular matrix is a prerequisite for healthy adipose tissue expansion, whereas fibrosis is a key feature of adipose dysfunction and inflammation. However, very little is known about the direct effects of impaired cell-matrix interaction in adipocyte function and insulin sensitivity. Using adipose selective deletion of β1 integrin (Itgb1^adipo-cre^) and Kindlin-2 (Kind2^adipo-cre^), we demonstrate here that active β1 and β3 integrins directly interact with the insulin receptor to regulate white adipocyte insulin action and systemic metabolism. Consequently, loss of adipose integrin activity, similar to loss of adipose insulin receptors, results in lipodystrophy and systemic insulin resistance. Conversely, we find that brown adipose tissue of Kind2^adipo-cre^ and Itgb1^adipo-cre^ mice is chronically hyperactivated, and has increased substrate delivery, reduced endothelial basement membrane thickness, and increased endothelial vesicular transport. Thus, we establish integrin-extracellular matrix interactions as key regulators of white and brown adipose tissue function and whole body metabolism.

## Introduction

Adipose tissue is essential to establish and maintain a functional metabolism (Luo, 2016). A prerequisite for proper adipose tissue function during development and weight gain is its ability to grow through de novo adipogenesis (hyperplasia) or expansion of preexisting adipocytes (hypertrophy). Both processes require reorganization of the extracellular matrix (ECM) to provide ample space for adipocytes and proper vascularization and innervation of the tissue. Thus, impaired adipose tissue expandability results in local hypoxia, tissue inflammation, insulin resistance and adipocyte death (Schoettl, Fischer et al., 2018, Sun, Kusminski et al., 2011, Sun, Tordjman et al., 2013), resulting in lipid spill over to the liver and skeletal muscle and the development of systemic insulin resistance and the metabolic syndrome (Boucher J, 2014). Integrins are a 24 member family of heterodimeric cell surface receptors composed of an α and a β subunit, essential to organize the ECM, facilitate anchorage of cells to the ECM and to initiate diverse intracellular signaling (Yoshikazu Takada, 2007). Moreover, altered integrin signaling, upon diet-induced obesity, associates with insulin resistance (Chloé C. Féral, 2008, Kang L, 2011, Williams, Kang et al., 2015). Integrins reside at the cell surface in a bend low-affinity state and require inside-out activation to shift to a high-affinity confirmation enabling binding to ECM components. This activation process is facilitated by binding of intracellular adaptor proteins of the Talin and Kindlin family (Calderwood, 2013, Theodosiou M, 2016). Kindlin-2 is ubiquitously expressed (Ussar, 2006) and essential for the activation of β1 and β3 containing integrins, in absence of other Kindlins (Harburger, Bouaouina et al., 2009, Kloeker, Major et al., 2004, Ma, Qin et al., 2008, Montanez, Ussar et al., 2008, Shi, Ma et al., 2007). There are no data on the role of talins or integrins in adipose tissue, but adipose specific loss of Kindlin-2 (Gao, Guo et al., 2019), similar to the loss of the key outside-in signal mediator focal adhesion kinase (Luk, Shi et al., 2017), results in lipodystrophy due to adipocyte apoptosis. Integrin signaling is very diverse and the signaling overlap between integrins and the insulin receptor suggest a potential effect of integrin signaling on insulin action. Indeed, α5β1 integrin activation enhanced basal insulin receptor phosphorylation as well as insulin stimulated insulin receptor substrate-1 (IRS-1) phosphorylation and recruitment of PI 3-kinase to IRS-1 (Guilherme A, 1998). Moreover, αvβ3 integrins and insulin signaling interact with each other (Schneller M, 1997, Vuori K, 1994). However, the *in vivo* relevance of this signal interaction, as well as the mechanistic details remain unknown. In contrast, previous work showed that β1 integrins can directly interact with the IGF1R in skeletal muscle to regulate IGF-1 mediated skeletal muscle growth (Wang, 2008). Thus, while multitudes of studies have investigated adipose tissue expansion and ECM remodeling and its metabolic consequences, the role of adipocyte cell surface receptors has not been addressed.

Here, we show that β1 and β3 integrins are expressed in adipocytes and their expression in subcutaneous and visceral fat correlates with obesity and insulin sensitivity in mice and humans. Using adipose specific knockouts of β1 integrin and Kindlin-2 we show that complete loss of integrin activity through deletion of Kindlin-2 is necessary to strongly impair white adipose tissue function, while loss of β1 integrin appears to be compensated by β3 integrins. Mechanistically, loss of Kindlin-2 impairs insulin action as active β1 and β3 integrins interact with the insulin receptor to regulate its activity. Moreover we demonstrate that loss Kindlin-2 results in hyperactivated brown adipose tissue, which is due to a completely new mechanism based on increased substrate delivery through the endothelium to brown adipocytes a proves that largely depends on active β1 integrins.

## Results

### Expression of β1 and β3 integrins in adipose tissue positively correlates with obesity and insulin resistance in humans

To study the relationship between body fat content and systemic insulin sensitivity, we analyzed gene expression of β1 and β3 integrins in subcutaneous (SAT) and visceral adipose tissue (VAT) in 123 subjects with a range of BMI. We found a positive correlation between expression of both β1 and β3 integrins with body fat content in VAT (**Fig. 1A)** and SAT (**Fig. EV1A**).Furthermore, expression of both integrins in VAT, but not in SAT, positively correlated with insulin resistance, estimated by the homeostasis model assessment of insulin resistance (HOMA-IR), which we adjusted for body fat percentage in a subgroup of non-diabetic subjects (n=115) (**Fig. 1B and Fig. EV1B**). Next, we analyzed protein levels of both β integrins, FAK, and Kindlin-2 in subcutaneous (SCF) and perigonadal (PGF) adipose tissue of mice fed either a chow (CD) or high fat diet (HFD) for 14 weeks. Protein levels of β1 and β3 integrins were significantly increased upon HFD feeding (**Fig. 1C**), albeit β1 integrin levels showed a greater increase compared to β3 integrins. Kindlin-2, p-FAK (Tyr387), and FAK were increased in SCF and PGF upon HFD feeding (**Fig. 1C**), as previously shown for FAK (Luk et al., 2017). Thus, integrins and integrin activity appear to be upregulated upon obesity in adipose tissues of mice and humans.

**Figure 1:**
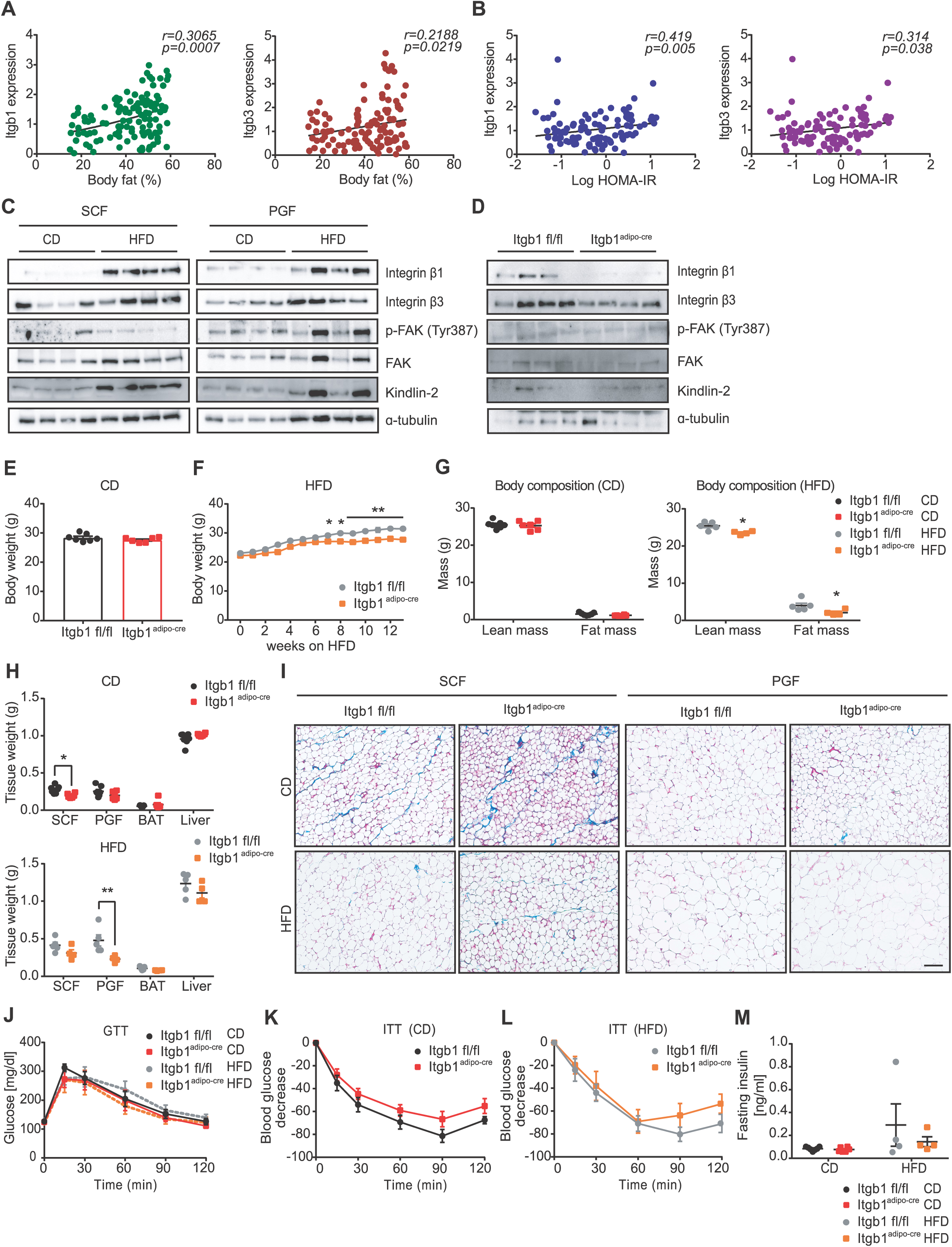
Expression of β1 and β3 integrins in adipose tissue are positively correlated with obesity and insulin resistance in humans and loss of β1 integrin in adipose tissue moderately impairs HFD induced-obesity. (**A**) Pearson correlation between the both beta 1 (β1) and beta 3 (β3) integrins in visceral adipose tissue (VAT) and body fat (%) in subjects of a wide distribution of BMI (n=123). (**B**) Partial correlation, adjusted by body fat (%), between both β1 and β3 integrins in VAT and insulin resistance measured by HOMA-IR in subjects with normal glucose metabolism (n=115). (**C**) Representative Western blot images for integrin β1, integrin β3, phospho(Tyr397)- and total FAK, Kindlin-2, and α-tubulin in subcutaneous (SCF) and perigonadal adipose tissue (PGF) in both chow (CD) and high-fat diet (HFD) fed wild-type mice (n=4). (**D**) Representative western blot images for integrin β1, integrin β3, phospho(Tyr397)- and total FAK, Kindlin-2, and α-tubulin in isolated adipocytes of SCF from integrin beta 1 flox mice (Itgb1 fl/fl) and Itgb1 flox; adiponectin-cre (Itgb1^adipo-cre^) mice fed CD for 13 weeks (n=4). (**E**) Body weight of Itgb1 fl/fl and Itgb1^adipo-cre^ mice fed CD for 13 weeks (n=7-6). (**F**) Body weight development of Itgb1 fl/fl and Itgb1^adipo-cre^ mice fed HFD for 14 weeks (n=5-4). (**G**) Body composition of Itgb1 fl/fl and Itgb1^adipo-cre^ mice fed CD (n=7-6) and HFD fed mice for 14 weeks (n=5-4). (**H**) Tissue weights of SCF, PGF, brown adipose tissue (BAT), and liver of Itgb1 fl/fl and Itgb1^adipo-cre^ mice fed CD (n=7-6) and HFD for 14 weeks (5-4). (**I**) Representative H&E and Masson trichrome staining of SCF and perigonadal adipose tissue (PGF) sections from Itgb1 fl/fl and Itgb1^adipo-cre^ mice fed CD and HFD for 14 weeks (scale bar represents 100 µm). (**J**) Plasma glucose concentration during intraperitoneal glucose tolerance test (2 g/kg) in Itgb1 fl/fl and Itgb1^adipo-cre^ mice fed CD (n=7-6) and HFD for 14 weeks (n=5-4). (**K**) Plasma glucose concentration during intraperitoneal insulin tolerance test (0.75 IU/kg) in Itgb1 fl/fl and Itgb1^adipo-cre^ mice fed CD for 14 weeks (n=7-6). (**L**) Plasma glucose concentration during intraperitoneal insulin tolerance test (1.25 IU/kg) in Itgb1 fl/fl and Itgb1^adipo-cre^ mice fed HFD for 14 weeks (n=5-4). (**M**) Fasting insulin levels (ng/ml) in Itgb1 fl/fl and Itgb1^adipo-cre^ mice fed CD (n=7-6) and HFD for 14 weeks (n=5-4). Data are shown as mean ± SEM. Statistics were calculated using ordinary two-way ANOVA with Tukey’s multiple comparison post-hoc test (**p<0.01, *p<0.05).

### Adipose-specific loss of β1 integrin moderately impairs HFD induced weight gain

β1 integrin containing integrins are the largest group of integrins binding to a diverse set of ECM components and thought to be the predominant integrins in adipocytes (Heemskerk, Mattheij et al., 2013, Hynes, 2002). Moreover, correlation with body fat content in mice and humans was stronger for β1 than β3 integrins. Thus, to explore the role of integrins in adipose tissue we generated a conditional knockout of β1 integrin in adipose tissue by crossing β1 integrin floxed mice with adiponectin-cre C57BL/6J mice (Itgb1^fl/fl^ are WT; and Itgb1^adipo-cre^ are conditional knockout). β1 integrin protein levels were reduced in total SCF lysates (**Fig. EV1C**) and undetectable in isolated subcutaneous adipocytes (**Fig.1D**), confirming efficient deletion of β1 integrin specifically in adipocytes. Loss of β1 integrin did not alter body composition in 13 week old CD fed mice (**Fig. 1E**), but significantly reduced body weight after 8 weeks on HFD (**Fig. 1F**), due to reduced fat and adipose tissue mass (**Fig. 1G** and **H**). Both, adipose selective depletion of Kindlin-2 and FAK, bluntin inside-out and outside-in integrin signaling, respectively, resulted in increased adipocyte death and loss of Kindlin-2 was reported to impair adipogenesis (Gao et al., 2019, Luk et al., 2017). However, we did not observe differences in *Pparg* and *Adiponectin* gene expression measured in primary subcutaneous adipocytes, indicating that adipogenesis per se is not impaired (**Fig. EV1D**). Moreover, H&E and Masson Trichrome stainings did not indicate any abnormal ECM deposition or immune cell infiltrations in either SCF or PGF in CD and HFD fed mice (**Fig. 1I**). Consequently, the slightly reduced fat mass did not affect glucose and insulin tolerance (**Fig. 1J-L**), fasting insulin (**Fig. 1M**), and glucose levels (**Fig. EV1E**) upon either CD or HFD feeding. Taken together, adipocyte specific loss of β1 integrin did not significantly impair adipose tissue function and whole body metabolism. However, loss of β1 integrin also did not blunt overall integrin signaling as monitored by p-FAK (**Fig. 1D**), strongly suggesting compensation by other integrins, most likely β3 integrins. To study the role of integrin activity in adipose tissue more generally, we generated adipose-specific loss of Kindlin-2, which regulates activation of β1 and β3 integrins.

### Adipose-specific loss of Kindlin-2 blunts integrin activity, causing age-dependent lipodystrophy

We created a conditional knockout of Kindlin-2 in adipose tissue in C57BL/6J mice, breeding adiponectin-cre with Kind2^fl/fl^ mice (Kind2 fl/fl are WT; and Kind2^adipo-cre^ are conditional KO). Kindlin-2 protein levels were strongly reduced in SCF, PGF and BAT upon CD or HFD feeding (**Fig. 2A** and **EV2A**). Loss of Kindlin-2 completely blunted FAK (Tyr 397) phosphorylation (**Fig. 2A**) and Rac1 activity (**Fig. 2B**). Thus, loss of Kindlin-2 indeed inactivates all relevant integrins in adipocytes and p-FAK as well as Rac1 activity can be used to assess integrin activity in adipocytes. Unexpectedly, loss of Kindlin-2 also significantly reduced β1 integrin protein levels, but had no effect on β3 integrins (**Fig. 2A**).

**Figure 2:**
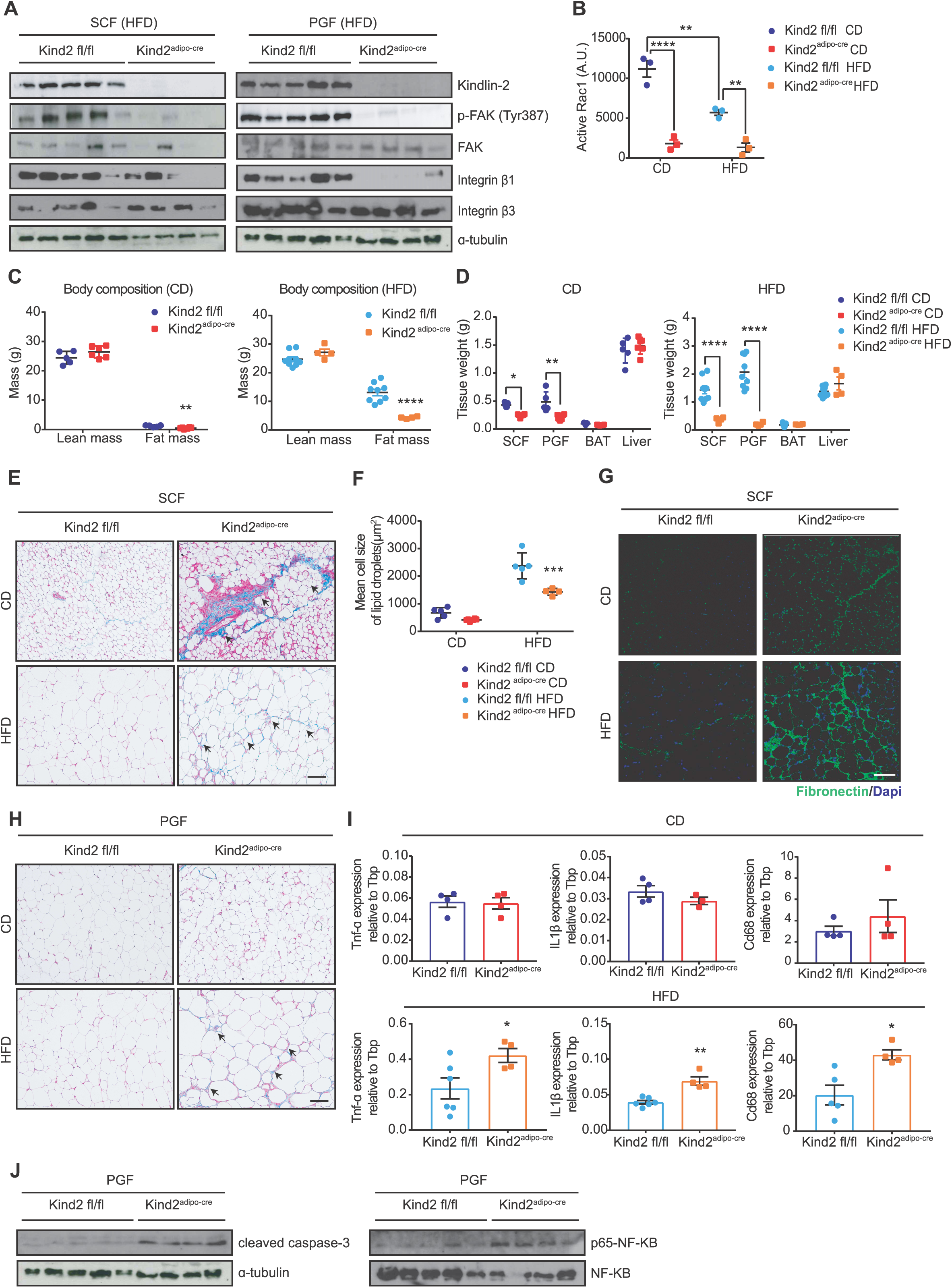
Adipose-specific loss of Kindlin-2 blunts integrin activity, causing age-dependent lipodystrophy. (**A**) Representative western blot images for Kindlin-2, phospho(Tyr397)- and total FAK, integrin beta 1 (β1), integrin beta 3 (β3) and α-tubulin in subcutaneous adipose tissue (SCF) and perigonadal adipose tissue (PGF) from Kindlin-2 flox (Kind2 fl/fl) and Kindlin-2 flox; adiponectin-cre (Kind2^adipo-cre^) mice fed HFD mice for 14 weeks. (**B**) Rac1 activation (small GTPases) in SCF of Kindlin-2 fl/fl and Kind2^adipo-cre^ mice fed CD and HFD fed mice for 14 weeks (n=3). (**C**) Body composition of Kind2 fl/fl and Kind2^adipo-cre^ mice fed CD (n=5-6) or HFD for 14 weeks (n=9-4). (**D**) Tissue weights of SCF, PGF, BAT, and liver of Kind2 fl/fl and Kind2^adipo-cre^ mice fed CD (n=5-6) or HFD for 14 weeks (n=9-4). (**E**) Representative H&E and Masson trichrome staining of SCF sections from Kind2 fl/fl and Kind2^adipo-cre^ mice fed CD and HFD for 14 weeks. Arrow bars represent signs of inflammation (scale bar represents 100 µm). (**F**) Mean cell size quantification of lipid droplets (µm^2^) in SCF from Kind2 fl/fl and Kind2^adipo-cre^ mice fed CD (n=5-5) and HFD for 14 weeks (n=5-4). (**G**) Immunofluorescence staining of fibronectin (green) and DAPI (blue) in SCF sections from Kind2 fl/fl and Kind2^adipo-cre^ mice fed CD and HFD for 14 weeks (scale bar represents 100 µm). (**H**) Representative H&E and Masson trichrome staining of PGF sections from Kind2 fl/fl and Kind2^adipo-cre^ mice fed CD and HFD for 14 weeks. Arrow bars represent signs of inflammation (scale bar represents 100 µm). (**I**) Relative gene expression of *Tnf-α, II1-β*, and *Cd-68* in PGF from Kind2 fl/fl and Kind2^adipo-cre^ mice fed CD (n=4) and HFD for 14 weeks (n=6-4). (**J**) Representative western blot images for cleaved caspase-3 and α-tubulin; and for phospho-(p65)-NF-κB and total NF-κB in PGF from Kind2 fl/fl and Kind2^adipo-cre^ mice fed HFD for 14 weeks (n=5-4). Data are shown as mean ± SEM. Statistics were calculated using two-way ANOVA with Tukey’s multiple comparison post-hoc test (***p<0.001, **p<0.01, *p<0.05).

Loss of Kindlin-2/ integrin activity reduced adipose tissue mass already at 10 weeks of age in CD fed mice (**Fig. EV2B**), before changes in overall body weight were detected (**Fig. EV2C**). Following 14 weeks of CD or HFD feeding, Kind2^adipo-cre^ mice showed a significant decrease in fat mass and adipose tissue weights (**Fig. 2C** and **D**), which translated to a decrease in overall body weight of HFD fed mice (**Fig. EV2D** and **EV2E**).

Consistent with the reduced fat mass, circulating leptin levels were markedly decreased in Kind2^adipo-cre^ mice under HFD, but not in CD fed mice (**Fig. EV2F**). However, free-fatty acids (FFA) in serum did not significantly differ between groups (**Fig. EV2G**). Importantly, we observed an identical phenotype in female Kind2^adipo-cre^ mice with regard to body weight development (**Fig. EV2H**), fat mass reduction (**Fig. EV2I**), and the decrease in fat pad weights (**Fig. EV2J**), resulting in an overall identical appearance upon HFD feeding (**Fig. EV2K**). Thus, integrin inactivation in adipose tissue, mediated by loss of Kindlin-2, blunts HFD-induced obesity by reducing adipose tissue mass in an age-dependent manner, suggesting an impact on adipose tissue expansion.

Both CD and HFD fed Kind2^adipo-cre^ mice showed reduced WAT mass, which could be protective from obesity associated metabolic complications, such as systemic insulin resistance and impaired glucose homeostasis. However, similar to lipodystrophy, loss of Kindlin-2 in adipose tissue resulted in adipose tissue fibrosis in CD and HFD fed mice (**Fig. 2E-H**), as well as increased inflammation and apoptosis upon HFD feeding (**Fig. 2I-J**). Kindlin-2 deficient primary subcutaneous adipocytes also showed increased *Tnf-α* expression, whereas expression of *Il1-β* or *Pparg*, was not statistically different (**Fig. EV2L**). Normal *Pparg* expression indicated that adipogenesis per se is not altered in adipocytes upon adiponectin-cre mediated deletion of Kindlin-2, which was also confirmed by differentiating primary brown adipocytes *in vitro* (**Fig. EV2M** and **N**).

### Active β1 and β3 integrins interact with the insulin receptor to modulate insulin action

The overall adipose tissue phenotype of Kind2^adipo-cre^ mice showed many similarities to the adipose tissue knockout of the insulin receptor (Qiang, 2016). Furthermore, previous studies suggested an interaction between integrin and insulin signaling (Guilherme A, 1998). Based on the data by Wang et al. (Wang, 2008), we tested the hypothesis that active β1 and β3 integrins can directly interact and modulate insulin receptor function. Co-immunoprecipitations (Co-IP) confirmed an interaction between β1 and β3 integrins with the insulin receptor in adipocytes and adipose tissue, respectively. This interaction was significantly reduced upon inactivation of the receptors by knockout of Kindlin-2 (**Fig. 3A**). Moreover, we found that protein levels of the insulin receptor were reduced in Kindlin-2 knockout mice (**Fig. 3B**). Insulin stimulation of primary perigonadal adipocytes from eight-week-old CD fed mice revealed strongly impaired Akt Ser473 and Thr308 phosphorylation (**Fig. 3C and Fig. EV3A**). Furthermore, insulin stimulated glucose uptake was significantly reduced (**Fig. 3D**). All together, these data demonstrated that active integrins interact with the insulin receptor and inactivation of integrins impairs insulin action. Interestingly, adipose selective knockout of mTORC1, through conditional ablation of raptor, resulted in a very similar lipodystrophic phenotype (Lee, Tang et al., 2016). *In vivo* insulin stimulation showed reduced Akt and p70-S6K phosphorylation in adipose tissue after 10 minutes in HFD fed Kind2^adipo-cre^ mice (**Fig. 3E**). Interestingly, while Akt phosphorylation was also reduced in BAT, although to a lesser extent than in WAT following intravenous insulin administration, we did not observe changes in phospho-p70-S6K (**Fig. 3E**).

**Figure 3:**
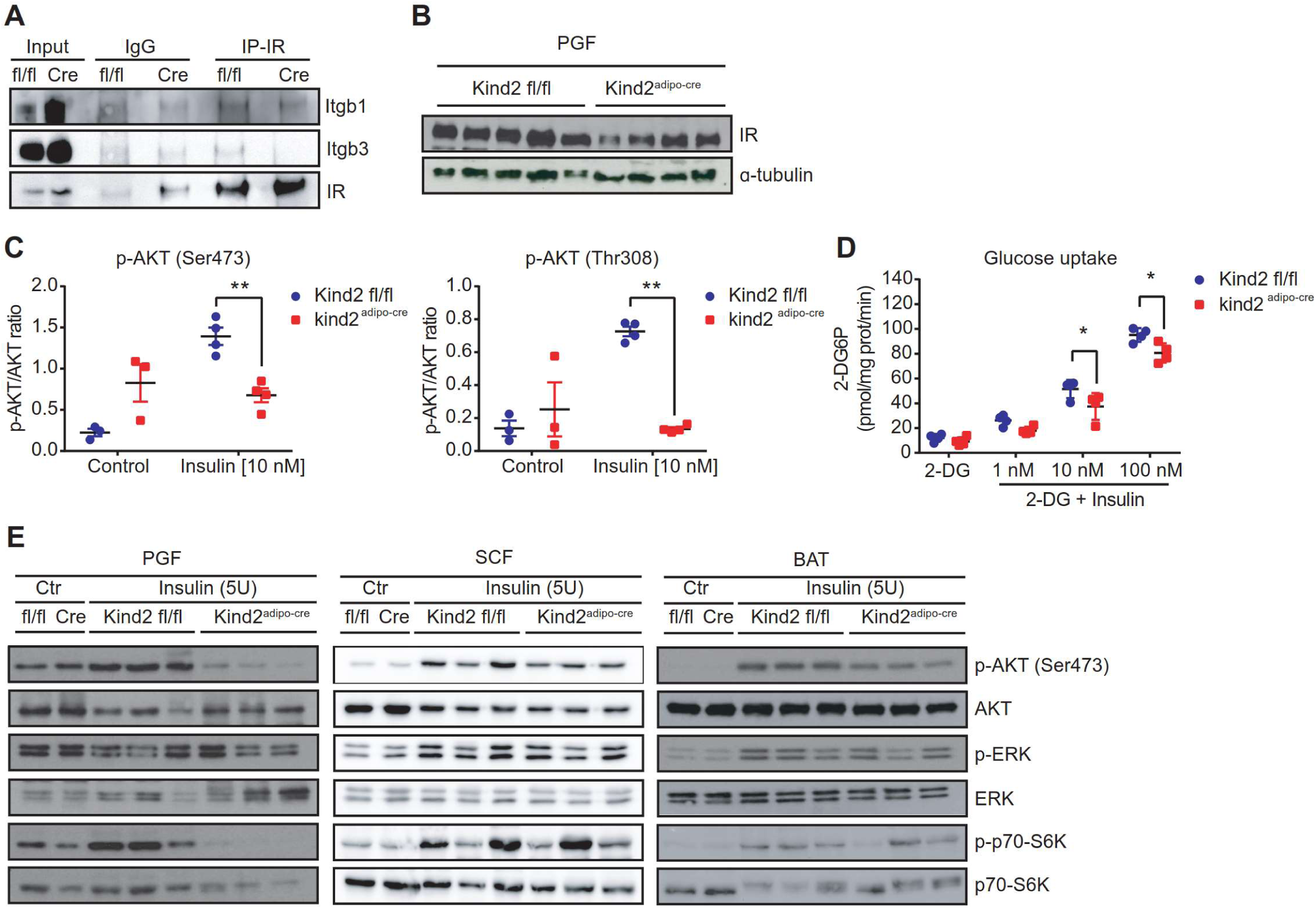
Active β1 and β3 integrins interact with the insulin receptor to modulate insulin action. (**A**) Representative western blot for co-immunoprecipitation of insulin receptor together with both β1 and β3 integrins in perigonadal adipose tissue (PGF) from Kind2 fl/fl and Kind2^adipo-cre^ mice fed CD for 8 weeks. (**B**) Representative western blot images for insulin receptor and α-tubulin in PGF from Kind2 fl/fl and Kind2^adipo-cre^ mice fed HFD for 14 weeks (n=5-4). (**C**) Western blot quantification for phospho-AKT (Ser473)/AKT and phospho-AKT (Thr308)/AKT in primary adipocytes isolated from PGF of Kindlin-2 fl/fl and Kind2^adipo-cre^ mice fed CD for 8 weeks in the presence or absence (control) of insulin [10 nM] for 10 min (n=3). (**D**) Ratio of glucose uptake (pmol/mg prot/min) in primary adipocytes isolated from PGF of Kindlin-2 fl/fl and Kind2^adipo-cre^ mice fed CD for 8 weeks in the presence or absence of insulin [10 nM] for 10 min (n=4). 2-DG is 2-deoxyglucose. (**E**) Representative western blot images for phospho-AKT (Ser473), AKT, phospho-ERK, ERK, phospho-p70-S6K and S6K in PGF, SCF and brown adipose tissue (BAT) in the presence or absence (ctr=control, saline) of insulin from *in vivo* insulin stimulation in Kind2 fl/fl and Kind2^adipo-cre^ mice fed HFD for 14 weeks (n=3). Data are shown as mean ± SEM. Statistics were calculated using two-way ANOVA with Tukey’s multiple comparison post-hoc test (**p<0.01, *p<0.05).

### Loss of Kindlin-2 in adipose tissue causes systemic insulin resistance without hepatosteatosis and hepatic insulin resistance

Loss of Kindlin-2 in adipocytes resulted in a lipodystrophic phenotype. In line with this Kind2^adipo-cre^ mice developed insulin resistance upon 14 weeks of HFD feeding (**Fig. 4A-B**). Glucose tolerance was not impaired in Kind2^adipo-cre^ mice (**Fig. 4C**), due to a compensatory increase in insulin secretion as evidenced by random fed hyperinsulinemia (**Fig. 4D**). Interestingly, however, we did not observe increased liver triglycerides (**Fig. 4E-G**). The same phenotype was also observed when high fat feeding was extended to 20 weeks (**Fig. EV4A-G**). Moreover, ERK phosphorylation was higher in the liver of Kind2^adipo-cre^ mice compared to controls when fed a HFD (**Fig. 4H**), indicating potentially increased hepatic insulin sensitivity, whereas the opposite would be expected in a lipodystrophic mouse. In line with this, we also observed significantly lower *Pepck* expression in the liver (**Fig. 4I**) and reduced fasting glycemia (**Fig. 4J**) after 14 weeks on HFD feeding. All these data indicated increased hepatic insulin sensitivity. *In vivo* insulin stimulation confirmed increased Akt phosphorylation at Ser473 in the liver from Kind2^adipo-cre^ mice compared to littermate controls (**Fig. 4K**). Thus, loss of Kindlin-2/ integrin activity in adipose tissue results in an atypical lipodystrophic phenotype with systemic insulin resistance that preserves liver insulin sensitivity.

**Figure 4.**
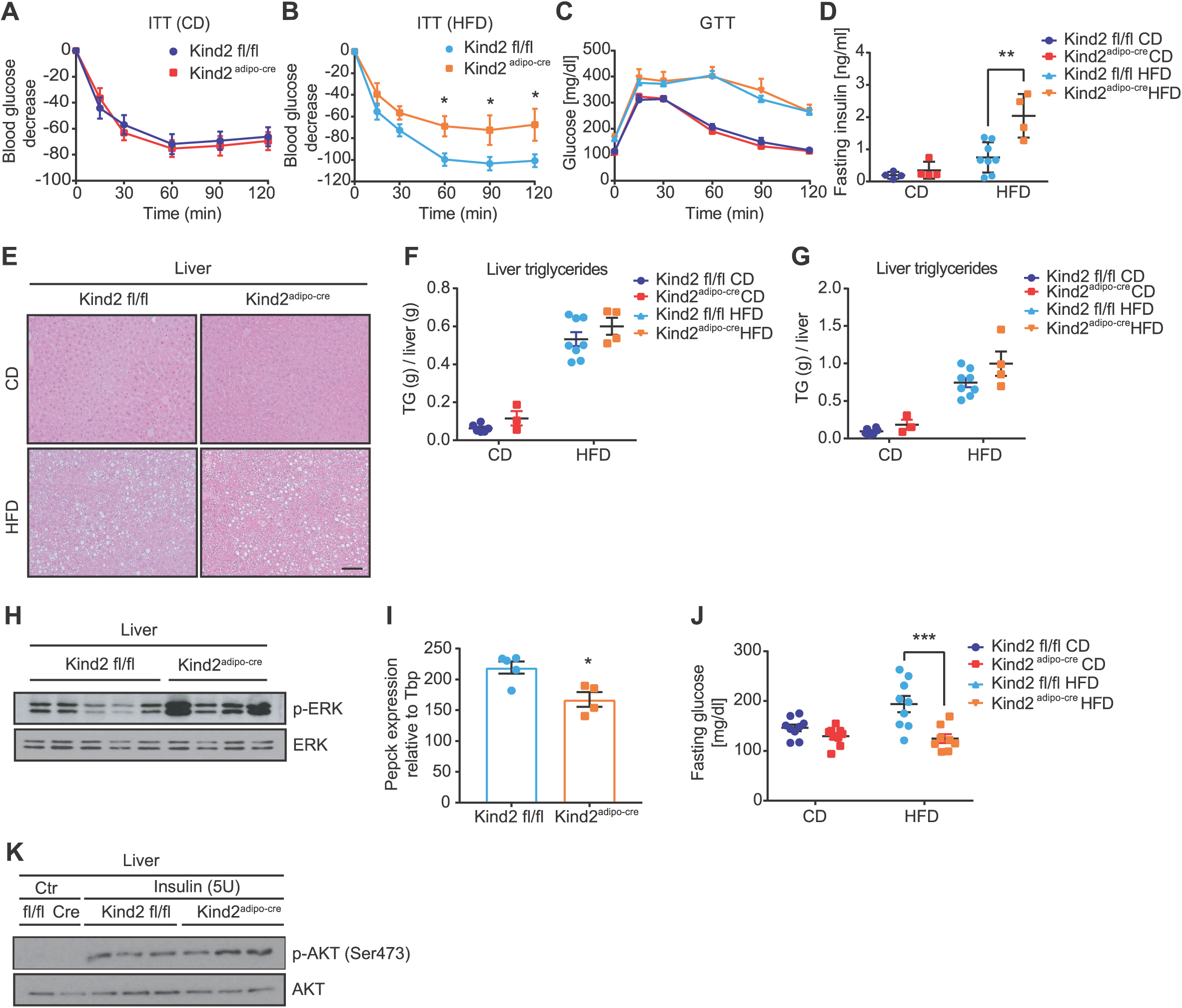
Loss of Kindlin-2 in adipose tissue causes systemic insulin resistance without hepatosteatosis and hepatic insulin resistance. (**A**) Plasma glucose concentration during intraperitoneal insulin tolerance test (0.75 IU/kg) in Kindlin-2 flox (Kind2 fl/fl) and Kindlin-2 flox; adiponectin-cre (Kind2^adipo-cre^) mice fed CD (n=7) 14 weeks (n=9-4). (**B**) Plasma glucose concentration during intraperitoneal insulin tolerance test (1.25 IU/kg) in Kind2 fl/fl and Kind2^adipo-cre^ mice fed HFD for 14 weeks (n=9-4). (**C**) Plasma glucose concentration during intraperitoneal glucose tolerance test (2 g/kg) in Kind2 fl/fl and Kind2^adipo-cre^ mice fed CD (n=7) and HFD for 14 weeks (n=9-4). (**D**) Fasting insulin levels (ng/ml) in Kind2 fl/fl and Kind2^adipo-cre^ mice fed CD (n=4) and HFD for 14 weeks (n=8-4). (**E**) Representative H&E staining of liver sections from Kind2 fl/fl and Kind2^adipo-cre^ mice fed CD and HFD for 14 weeks (scale bar represents 100 µm). (**F**) Triglyceride content in liver (g/g) in Kind2 fl/fl and Kind2^adipo-cre^ mice fed CD (n=4-3) and HFD for 14 weeks (n=8-4). (**G**) Total triglyceride content (g) per whole liver in Kind2 fl/fl and Kind2^adipo-cre^ mice fed CD (n=4-3) and HFD for 14 weeks (n=8-4). (**H**) Representative western blot images for phospho-ERK and ERK in liver from Kindlin-2 fl/fl and Kind2^adipo-cre^ mice fed HFD for 14 weeks (n=5-4). (**I**) Relative gene expression relative to *Tbp* of *Pepck* in liver from Kind2 fl/fl and Kind2^adipo-cre^ mice fed HFD for 14 weeks (n=5-4). (**J**) Fasting glucose (mg/dl) in Kind2 fl/fl and Kind2^adipo-cre^ mice fed CD (n=9) and HFD for 14 weeks (n=9-8). (**K**) Representative western blot images for phospho-AKT (Ser473) and AKT in liver from Kind2 fl/fl and Kind2^adipo-cre^ mice fed CD for 13 weeks in the presence or absence (ctr=control, saline) of insulin (*in vivo* insulin stimulation) (n=3). Data are shown as mean ± SEM. Statistics were calculated using two-way ANOVA with Tukey’s multiple comparison post-hoc test (***p<0.001, **p<0.01, *p<0.05).

### Loss of Kindlin-2 increases BAT activity independent from diet and housing temperature

The absence of hepatosteatosis despite loss of adipose tissue mass suggested reduced food intake or increased energy expenditure. Assessment of food and water intake, as well as the respiratory exchange ratio, using metabolic cages, did not reveal any differences between control and Kind2^adipo-cre^ mice fed either chow or HF diet (**Fig. EV5A-C**). However, energy expenditure (EE) of Kind2^adipo-cre^ mice was increased in HFD but not CD fed mice (**Fig. 5A**), with a trend to higher UCP-1 protein levels upon HFD feeding (**Fig. 5B**), and decreased lipid droplet size in Kind2^adipo-cre^ mice fed either CD or HFD (**Fig. 5C-D**). Moreover, fecal energy content of HFD fed Kind2^adipo-cre^ mice was significantly higher compared to controls, whereas we did not observe differences in CD fed animals (**Fig. EV3B**), phenocopying loss of raptor in adipose tissue (Lee et al., 2016). Thus, a combination of increased brown fat activity together with elevated energy excretion through the feces appears to explain the absence of hepatosteatosis and hepatic insulin resistance in context of lipodystrophy of Kind2^adipo-cre^ mice. To further characterize the effect of adipose tissue loss of Kindlin-2 on BAT activity, HFD fed Kind2^adipo-cre^ and littermate control mice were housed at 4 °C for 48 h in metabolic cages. Kind2^adipo-cre^ mice showed increased EE (**Fig. 5E**), and elevated food intake (**Fig. 5F**) and RER (**Fig. 5G**). Cold exposure for 48 hours induces the formation of brown-like adipocytes, called “beige” or “brite” (brown-in-white) (Wu, Bostrom et al., 2012), within white adipose tissue depots. However, we did not observe significant differences in *Ucp1* and *Prdm16* expression in SCF (**Fig. EV5D**), albeit there was a clear trend for elevated Ucp1 expression in Kind2^adipo-cre^ mice. Similarly, we did not observe differences in *Ucp1* or *Prdm16* expression in BAT after chronic cold exposure (**Fig. EV5D**).

**Figure 5:**
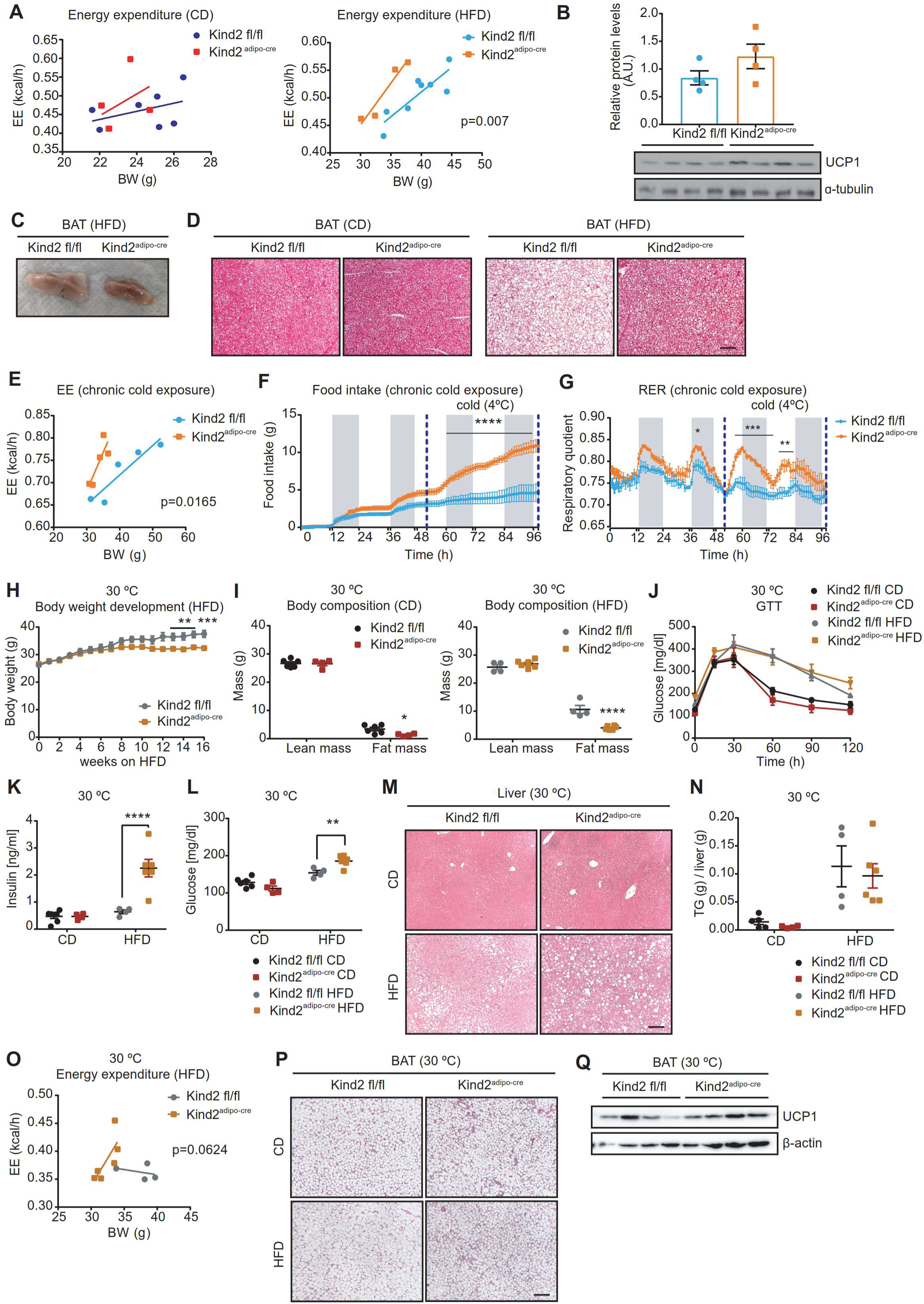
Loss of Kindlin-2 increases BAT activity independent from diet and housing temperature. (**A**) Body-weight (g) correlated to total energy expenditure (kcal/h) in Kindlin-2 flox (Kind2 fl/fl) and Kindlin-2 flox; adiponectin-cre (Kind2^adipo-cre^) mice fed CD (n=7-4) and HFD for 13 weeks (n=8-4). (**B**) Representative western blot images for UCP1 and α-tubulin in brown adipose tissue (BAT) from in Kind2 fl/fl and Kind2^adipo-cre^ mice fed HFD for 14 weeks (n=4). (**C**) Photograph of BAT from Kind2 fl/fl and Kind2^adipo-cre^ mice fed HFD for 14 weeks. (**D**) Representative H&E staining of BAT sections from in Kind2 fl/fl and Kind2^adipo-cre^ mice fed CD and HFD for 14 weeks (scale bar represents 100 µm). (**E**) Body-weight (g) correlated to total energy expenditure (kcal/h) in Kind2 fl/fl and Kind2^adipo-cre^ mice fed HFD for 14 weeks during chronic cold exposure at 4°C for 48 hours (n=5-4). (**F**) Cumulative food intake (g) for 4 days in in Kind2 fl/fl and Kind2^adipo-cre^ mice fed HFD for 14 weeks. Blue lines represent chronic cold exposure at 4 °C for 48 hours (n=5-4). (**G**) RER in Kind2 fl/fl and Kind2^adipo-cre^ mice fed HFD for 14 weeks during chronic cold exposure at 4 °C for 48 hours (n=5-4). Blue lines represent chronic cold exposure at 4 °C for 48 hours (n=5-4). (**H**) Body weight development of in Kind2 fl/fl and Kind2^adipo-cre^ mice fed HFD for 16 weeks housed at thermoneutrality (30°C) (n=4-6). (**I**) Body composition represented as lean and fat mass weights (g) of in Kind2 fl/fl and Kind2^adipo-cre^ mice fed CD (n=6-4) and HFD (n=4-6) for 16 weeks housed at thermoneutrality (30°C). (**J**) Plasma glucose concentration during intraperitoneal glucose tolerance test (2 g/kg) in Kind2 fl/fl and Kind2^adipo-cre^ mice fed CD (n=6-4) and HFD (n=4-6) for 16 weeks housed at thermoneutrality (30°C). (**K**) Fasting insulin levels (ng/ml) in mice Kind2 fl/fl and Kind2^adipo-cre^ mice fed CD (n=6-4) and HFD (n=4-6) for 16 weeks housed at thermoneutrality (30°C). (**L**) Fasting glucose levels (mg/dl) in mice Kind2 fl/fl and Kind2^adipo-cre^ mice fed CD (n=6-4) and HFD (n=4-6) for 16 weeks housed at thermoneutrality (30°C). (**M**) Representative H&E staining of liver sections from Kind2 fl/fl and Kind2^adipo-cre^ mice fed CD and HFD for 16 weeks housed at thermoneutrality (30°C) (scale bar represents 100 µm). (**N**) Triglyceride content in liver (g/g) in Kind2 fl/fl and Kind2^adipo-cre^ mice fed CD (n=5-4) and HFD (n=4-6) for 16 weeks housed at thermoneutrality (30°C). (**O**) Body-weight (g) correlated to total energy expenditure (kcal/h) in Kind2 fl/fl and Kind2^adipo-cre^ mice fed HFD (n=4-6) for 16 weeks housed at thermoneutrality (30°C). (**P**) Representative H&E staining of BAT sections from Kind2 fl/fl and Kind2^adipo-cre^ mice fed CD and HFD for 16 weeks housed at thermoneutrality (30°C) (scale bar represents 100 µm). (**Q**) Representative western blot images for UCP1 and α-tubulin in BAT from Kind2 fl/fl and Kind2^adipo-cre^ mice fed HFD (n=3-4) for 16 weeks housed at thermoneutrality (30°C). Data are shown as mean ± SEM. Statistics were calculated using two-way ANOVA with Tukey’s multiple comparison post-hoc test (****p<0.0001, ***p<0.001, **p<0.01, *p<0.05).

We next tested if inactivation of BAT in Kind2^adipo-cre^ mice through a combination of HFD feeding and housing at thermoneutrality, as suggested previously by others (Cannon B, 2004, Clayton ZS, 2018, Cui X, 2016), could result in hepatosteatosis and the development of metabolic complications classically associated with lipodystrophy. Housing HFD fed Kind2^adipo-cre^ mice at 30 °C impaired diet induced weight gain and accumulation of fat mass (**Fig 5H** and **I**) to a similar extend as mice housed at room temperature (23 °C), while CD feeding did not result in body weight differences (**Fig. EV5E**), but reduced fat mass (**Fig. 5I**). Assessment of glucose and insulin tolerance, as well as fasting insulin levels largely confirmed the insulin resistant phenotype observed at room temperature (**Fig. 5J-K, Fig. EV5F-G**). However, unlike at room temperature, at thermoneutrality HFD-fed Kind2^adipo-cre^ mice showed increased fasting glycemia (**Fig. 5L**), similar ERK activation (**Fig. EV5H**) and *Pepck* gene expression in liver (**Fig. EV5I**), when compared to littermate controls. Thus, while not affecting the phenotype in white adipose tissue, housing at thermoneutrality resulted in increased hepatic insulin resistance of Kind2^adipo-^ cre mice compared to Kind2^adipo-cre^ mice housed at room temperature. Nevertheless, also at thermoneutrality we did not observe ectopic lipid accumulation in liver (**Fig. 5M-N)** or skeletal muscle **(Fig. EV5J**). Consistently, we observed increased energy expenditure in Kind2^adipo-cre^ HFD fed mice housed at 30 °C (**Fig. 5O**), a trend to increased UCP1 protein levels and decreased BAT lipid droplet size in both CD and HFD fed mice (**Fig. 5P-Q**). Similar to mice housed at room temperature, food intake (**Fig. EV5K**) and RER (**Fig. EV5L**) were comparable to control mice.

Hence, Kind2^adipo-cre^ mice displayed a previously undescribed hyperactive brown fat even upon thermoneutrality and HFD feeding, a situation with low sympathetic activity.

### Loss of β1 integrin activity in brown adipocytes increases vascular permeability in BAT

BAT is highly vascularized with every brown adipocyte in contact with capillaries, only separated by a basement membrane (Peirce V., 2016). Thus, changes in the vascularization or vascular permeability could deliver more substrate to brown adipocytes driving thermogenesis through substrate excess rather sympathetic control. Indeed, intravenous perfusion with Evan’s blue showed increased dye penetration into BAT of Kind2^adipo-cre^ mice (**Fig. 6A-B**), indicative of increased vascular permeability. A similar phenotype was also observed in SCF and PGF (**Fig. EV6A**). However, blood vessel density and morphology appeared normal in BAT of Kindlin-2 knockout mice (**Fig. 6C**). Electron microscopy (EM) revealed that loss Kindlin-2 reduced basement membrane thickness between the endothelium and brown adipocytes (**Fig. 6D-E and EV6C**) and increased endothelial vesicle density (**Fig. 6F and EV6D**), whereas endothelial overlap was unaltered (**Fig. 6G**), indicating increased trans-endothelial transport to brown adipocytes. Importantly, we observed a similar increase in Evan’s blue penetration in BAT of β1 integrin knockout mice (**Fig. 6A-B**), suggesting that the increased vascular permeability is mediated through loss of β1 integrin function in brown adipocytes. Similar to Kindlin-2 knockout mice, SCF and PGF of Itgb1^adipo-cre^ mice also had increased Evan’s blue penetration (**Fig. EV6B**).

**Figure 6:**
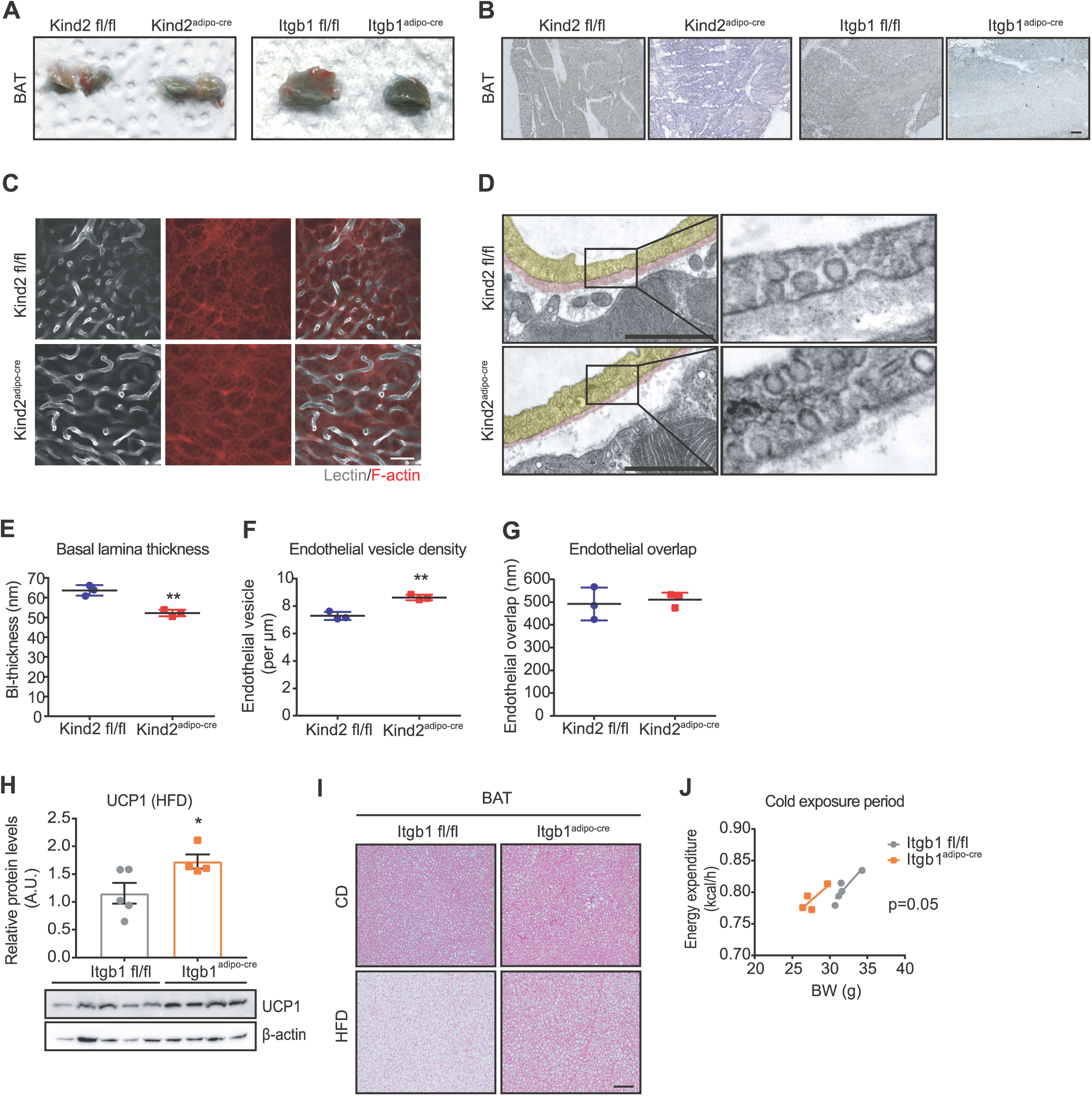
Loss of β1 integrin activity in brown adipocytes increases vascular permeability in BAT. (**A**) Photograph of brown adipose tissue (BAT) from Kindlin-2 flox (Kind2 fl/fl) and Kindlin-2 flox; adiponectin-cre (Kind2^adipo-cre^) mice fed CD for 8 weeks and integrin beta 1 flox mice (Itgb1 fl/fl) and Itgb1 flox; adiponectin-cre (Itgb1^adipo-cre^) mice fed CD for 8 weeks after Evans blue (0.05 % in PBS) perfusion and fixation with PFA (4 %). (**B**) Representative images from BAT cryo-sections (30 µm) after Evans blue perfusion and PFA fixation from Kind2 fl/fl and Kind2^adipo-cre^ and Itgb1 fl/fl and Itgb1^adipo-cre^ mice fed CD for 8 weeks (scale bar represents 100 µm). (**C**) Representative images from BAT of Kind2 fl/fl and Kind2^adipo-cre^ mice fed CD for 8 weeks after lectin perfusion and fixation with 4 % PFA. Lectin is represented in white color and F-actin in red (scale bar represents 25 µm). (**D**) Representative electron microscopy (EM) images showing adjacent parts of adipocytes in BAT from Kind2 fl/fl (WT) and Kind2^adipo-cre^ (KO) mice fed CD for 8 weeks (scale bar represents 1 µm). (**E**) Basal lamina thickness quantification (µm) measured by EM at 15 different points per vessels. Mean value is calculated per vessel and mouse in BAT from Kind2 fl/fl and Kind2^adipo-cre^ mice fed CD for 8 weeks (n=3). (**F**) Endothelial vesicle density (per µm) measured by EM in BAT from Kind2 fl/fl and Kind2^adipo-cre^ mice fed CD for 8 weeks (n=3). (**G**) Endothelial overlap (nm) measured by EM in BAT from Kind2 fl/fl and Kind2^adipo-cre^ mice fed CD for 8 weeks (n=3). (**H**) Representative western blot images for UCP1 and α-tubulin in BAT from Itgb1 fl/fl and Itgb1^adipo-cre^ mice fed HFD for 14 weeks (n=5-4). (**I**) Representative H&E staining of BAT sections from Itgb1 fl/fl and Itgb1^adipo-cre^ mice fed CD and HFD for 14 weeks (scale bar represents 100 µm). (**J**) Body-weight (g) correlated to energy expenditure (kcal/h) in Itgb1 fl/fl and Itgb1^adipo-cre^ mice fed HFD for 14 weeks during acute cold exposure at 4 °C for 5 hours (n=5-4). Data are shown as mean ± SEM. Statistics were calculated using two-way ANOVA with Tukey’s multiple comparison post-hoc test (**p<0.01, *p<0.05).

Thus, if increased vascular permeability drives increased energy expenditure, a similar effect should be observed in β1 integrin knockout mice. Indeed, *Ucp1* mRNA (**Fig. EV6F**) and protein levels (**Fig. 6H**) both on CD and on HFD were increased in β1 integrin knockout mice and lipid content in BAT reduced (**Fig. 6I**). Moreover, we observed increased energy expenditure upon acute cold exposure (**Fig. 6J**), without changes in food intake (**Fig. EV6G**) and RER (**Fig. EV6H**), supporting that the increased BAT activity in Kind2^adipo-cre^ mice is mediated by the regulation of β1 integrin orchestrated trans-endothelial transport.

## Discussion

Maintaining or restoring adipose tissue function is pivotal to prevent the progression from obesity towards insulin resistance and the metabolic syndrome. Here, we show that expression of β1 and β3 integrins in adipose tissues is positively correlated with obesity and insulin resistance in humans, and adipose-specific loss of β1 integrin in mice shows a moderate reduction in fat mass without metabolic impairments. General loss of integrin activity through adipose-specific knockout of Kindlin-2, however, dramatically limits adipose tissue expansion triggering fibrosis, inflammation and apoptosis in white, but not brown fat. This is very similar to the adipose tissue specific knockout of the insulin receptor (Qiang, 2016). Indeed, we find that Kindlin-2 deficient adipocytes, prior to the development of inflammation, are insulin resistant and that β1 and β3 integrins directly interact with the insulin receptor, which is significantly reduced upon loss of Kindlin-2 and thereby loss of integrin inside-out activation of integrins. Previous works by Michael Czech and Erkki Ruoslahti have already suggested an interaction of β1 and β3 integrins with insulin action, respectively, albeit the molecular details and physiological consequences remained unknown (Guilherme A, 1998, Vuori K, 1994).

Insulin and integrin signaling regulate various overlapping cellular pathways, of which many can contribute to the observed adipose dysfunction (Guilherme A, 1998). Among the various effectors, we find a reduction in mTORC1 activity in white but not brown adipocytes, which appears as the primary mediator of the observed lipodystrophy, inflammation, and fibrosis as adipose selective raptor knockout has an almost identical phenotype in white adipose tissue (Lee et al., 2016, Polak, Cybulski et al., 2008). mTORC1 integrates multiple upstream signals, including integrin and insulin receptor signaling (Bui, Rennhack et al., 2019, Dibble & Manning, 2013). However, loss of integrin activity was observed in both white and brown fat, but insulin resistance was restricted to WAT, as was the reduction in mTORC1 activity. Thus, the impaired mTORC1 activity observed in Kindlin-2 knockout mice appears to be due to impaired insulin rather than integrin signaling, further substantiating the significant role of integrins in the modulation of insulin action in white adipocytes. Importantly, the absence of adipocyte insulin resistance in β1 integrin deficient adipocytes suggests that β3 containing integrins can, at least, compensate for the loss of β1 integrins, essentially limiting the ECM components mediating these effects to RGD containing fibronectin and vitronectin (Charo IF, 1990, Horton, 1997). This raises the interesting possibility that changes in the composition of the ECM, such as during tissue expansion and inflammation, can directly and selectively impact on adipocyte insulin sensitivity. This is in line with a recent study showing that obesity-induced insulin resistance in mice can precede macrophage accumulation and inflammation in adipose tissue (Shimobayashi, Albert et al., 2018). Thus, integrin activity in mature white adipocytes appears to predominantly affect insulin action, which in some ways, is in contradiction to the conclusions drawn in a recent report on adipose selective knockout of Kindlin-2 (Gao et al., 2019). It is indeed difficult to conclude if the observed tissue fibrosis and apoptosis upon loss of Kindlin-2 are due to impaired integrin or insulin action, as both can mediate these processes. However, a dependency on insulin receptor action is more likely, as primary white adipocytes from young CD fed Kindlin-2 knockouts did not show increased expression of proinflammatory cytokines, suggesting an indirect effect. Furthermore, differentiation of primary preadipocytes did not show any impairment in adipogenesis or signs of anoikis, most likely due to the relatively late deletion of Kindlin-2 through cre expression under the control of the adiponectin promoter.

In contrast to WAT, loss of Kindlin-2/ integrin activity in BAT has limited effects on insulin sensitivity and mTORC1 activation. It will be interesting to determine if potential differences in the ECM between WAT and BAT could explain the minor role of integrins in regulating brown adipocyte insulin sensitivity or other signals compensate for the loss of integrins in the future. However, the fact that insulin/mTORC1 action is preserved in Kindlin-2 knockout mice helps to explain the systemic differences between adipose selective knockout of Kindlin-2 and raptor. Loss of raptor in adipocytes results in lipodystrophy and, if deleted using adiponectin-cre, insulin resistance, hepatosteatosis, and glucose intolerance. Loss of Kindlin-2 also results in systemic insulin resistance. However, we did not observe hepatosteatosis. In fact, the liver of Kind2^adipo-cre^ mice appeared protected from HFD induced steatosis and insulin resistance, suggesting that the energy that is not stored in white fat is either not taken in or expended. Fecal caloric content was increased in HFD fed Kind2^adipo-cre^ mice. However, the same was observed in conditional raptor knockout mice, which showed strong hepatosteatosis (Lee et al., 2016). Indirect calorimetry did not indicate differences in food intake or activity, but increased energy expenditure, especially when mice were fed a HFD. Surprisingly, however, energy expenditure of Kind2^adipo-cre^ mice, as well as UCP-1 protein levels in BAT were increased not only at room temperature and 4 °C, but also at thermoneutrality when combined with HFD feeding, a situation of low sympathetic activity, usually associated with inactivation of brown fat (Cui X, 2016).

This suggests that sympathetic signals are not the primary mediators of the increased BAT activity in adipose Kindlin-2 and β1 integrin knockouts. In addition to the dense innervation of BAT, every brown adipocyte is also in contact with blood vessels, only separated by a basement membrane (Peirce V., 2016). Fasting alters the endothelial basement membrane (BM) structure in BAT, resulting in increased transport into the intercellular space (Yudasaka, Yomogida et al., 2018) as well as increased glucose uptake into BAT of humans (Vrieze A, 2012). This suggests that changes in the endothelial BM in BAT can increase substrate delivery to brown adipocytes and thereby its activity. Integrins are important organizers of the basement membrane (Ojakian GK, 1994) and we find a reduction in the thickness of the endothelial BM in Kind2^adipo-cre^ mice. Moreover, we also find increased penetration of perfused Evans blue into BAT. However, this did not result in impaired endothelial cell-cell contacts, but rather increased endothelial vesicles, indicative of increased trans-endothelial transport. Therefore, we conclude that loss of integrin activation in brown adipocytes results in an impaired BM between brown adipocytes and endothelial cells causing increased trans-endothelial transport and more substrate delivery to brown adipocytes, thereby uncoupling brown adipocyte activity from sympathetic inputs. The absence of such a phenotype in adipose selective FAK knockouts (Luk et al., 2017) suggests that this is mediated through the role of integrins in organizing the ECM rather than transmitting signals intracellularly. This mechanism also explains the larger effect on energy expenditure upon HFD feeding of Kind-2^adipo-cre^ mice as, due to the inability of WAT to store excessive calories; more substrate is in the circulation. Furthermore, the hyperactivation of BAT also helps to explain the relative improvement in hepatic insulin sensitivity and the absence of ectopic lipid accumulation in skeletal muscle and liver in context of lipodystrophy. Interestingly, in contrast to WAT insulin sensitivity, the effect on endothelial transport seems to be largely dependent on β1 integrins, as loss of β1 integrin phenocopies the BAT phenotype of Kind2^adipo-cre^ mice.

Thus, our data establish integrin activity in adipose tissue as an important regulator of systemic metabolism, with a critical function in regulating white adipose tissue insulin sensitivity. The important metabolic role of BAT and the differential effects of loss of integrin function, as well as insulin action in brown versus white fat result in a complex metabolic phenotype. Loss of integrin activity in WAT causes age acquired lipodystrophy, associated with systemic insulin resistance, without impairing the liver due to a second function of integrins in BAT to regulate trans-endothelial substrate delivery by modulating BM thickness.

We observed a correlation of β1 and β3 integrin expression with obesity and metrics of insulin resistance in humans, suggesting a potentially conserved function between humans and mice. Large parts of human BAT are in an inactive state. However, as loss of β1 integrin activity in brown adipocytes uncouples BAT activity from sympathetic activity, it is tempting to speculate that similar to mice, impaired adipose integrin activity in humans also results in an atypical lipodystrophy with loss of white adipose tissue mass in the absence of ectopic lipid accumulation due to hyperactivation of BAT. White adipose tissue fibrosis and inflammation are hallmarks of adipose dysfunction and linked to insulin resistance and type 2 diabetes (Sun et al., 2013). However, our data suggest that changes in the composition and structure of the ECM surrounding white adipocytes can directly and profoundly change insulin sensitivity and directly contribute to the predisposition of patients to the development of metabolic complications (Kahn CR, 2019).

In summary, our study unravels novel concepts for the regulation of white adipocyte insulin receptor action and tissue expansion as well as brown fat activity by previously unrecognized distinct functions of active integrins in white and brown adipocytes.

## Material and Methods

### Mice

All mice were maintained in a conventional animal facility at constant ambient temperature of 22 ± 2°C, with 45-65% humidity and a 12h light-dark cycle in ventilated racks. Cages and water were supplemented with nesting material and were changed every week. All mice received a standard chow diet (Altromin 1314, Lage Germany) and water *ad libitum* until further dietary interventions or death. Adipose-selective loss of β1 integrin was generated by breeding Itgb1^flox/flox^ mice (exons 2 to 7 of Itgb1 gene were flanked by LoxP sites) (Potocnik AJ, 2000) with mice carrying Cre recombinase driven by the adiponectin promoter (Adipo-Cre) (Jackson Laboratories), both on a C57BL/6J background. Itgb1^fl/+^ heterozygous mice were bred to generate Adipo-Cre-positive littermate mice, and Adipo-Cre-positive males and Adipo-Cre-negative female mice were used for breeding. Male and female mice were used throughout the study, using as a control Adipo-Cre-negative floxed (Itgb1^fl/fl^) and the adipose-specific knockout Adipo-Cre-positive floxed (Itgb1^adipo-cre^). Kindlin-2 (Fermt2) floxed mice were obtained by breeding Fermt2tm1a mice (EUCOMM)Wtsi (IMPC, GSF-EPD0087_1_G04-1) with Flp-deleter mice to remove the stop cassette and subsequently mating with C57BL/6J mice to remove the Flp transgene. Kind2^flox/+^ mice were intercrossed to obtain Kind2^flox/flox^ mice. Adipose-selective loss of Kindlin-2 was generated by breeding Kind2^flox/flox^ mice (exons 5 and 6 of Fermt2 gene were flanked by LoxP sites) with mice carrying Cre recombinase driven by the adiponectin promoter (Adipo-Cre) (Jackson Laboratories), both on a C57BL/6J background. Male and female mice were used throughout the study, using as a control Adipo-Cre-negative floxed (Kind2^fl/fl^) and the adipose-specific knockout Adipo-Cre-positive floxed (Kind2^adipo-cre^). 58% High fat diet (HFD) was used for HFD feeding starting at 8 weeks-old (Research Diets D12331, New Brunswick (NJ), USA) for a duration of 14 weeks. Body weight was determined in at weekly intervals and body composition was measured by nuclear magnetic resonance measurements (EchoMRI ^®^LLC, Houston, USA) before the onset of the HFD feeding and prior to sacrificing the mice. For the thermoneutrality study, mice were housed at 30°C. For cold exposure, mice were singly housed at 4 °C for 2 days for chronic cold exposure and 4 °C for 5 hours for acute chronic cold exposure. Animal experiments were conducted in accordance with the German animal welfare law and performed with permission and in accordance with all relevant guidelines and regulations of the district government of Upper Bavaria (Bavaria, Germany), protocol number 55.2-1-54-2532-52-2016.

### Genotyping of mouse lines

Eartags were obtained from mice at the age of 3 weeks, and DNA was isolated by boiling the eartags for 30 min in 100 µl 50 mM NaOH at 95 °C (ThermoMixer C, Eppendorf). Afterwards, 10 µl 1 M Tris was added to normalize the pH. 1 µl of isolated genomic DNA was used for the genotyping PCR (Promega) using respective protocols.

### Energy metabolism studies

Body composition (fat and lean mass) was analyzed using a magnetic resonance whole-body composition analyzer (EchoMRI, Houston, TX). Assessment of energy intake, water intake, respiratory exchange quotient (RER), and energy expenditure and home-cage activity were performed using an indirect calorimetry system (TSE PhenoMaster, TSE Systems, Bad Homburg, Germany). Following 24 h of acclimatization, O_2_ consumption, and CO_2_ production were measured every 10 min for a total of 72 h. Linear regression was used to analyze the energy expenditure, with body weight as covariate. Fecal caloric content was measured from dried fecal pellets collected from single cages using a 6300 Oxygen Bomb Calorimeter (Parr Instrument Technology).

### Glucose and insulin tolerance tests

Glucose tolerance tests (GTT) and insulin tolerance tests (ITT) were carried out in mice fasted for 4 hours. 2 g/kg glucose (20 % Glucose solution; Braun) or 0.75/1.25 U insulin/ kg BW (Actrapaid® PenFill® Novo Nordisk) were injected intraperitoneally (i.p.) and glucose concentrations measured in blood collected from the tail before and 15, 30, 60, 90, and 120 min after the injection using a FreeStyle Freedom Lite glucometer (Abbott).

### Systemic insulin stimulation *in vivo*

Mice were fasted for 4 h, and recombinant human insulin (100 IU/mL, 50 μL; Humalin R) or saline was injected via the inferior vena cava after anesthesia with ketamine (100 mg/kg) and xylazine (5 mg/kg). The procedure was carried out under deep anesthesia and with the administration of metamizol (200 mg/kg) as analgesic. During the experiment, mice were placed on a heating pad to prevent hypothermia. At 10 min after the injection, SCF, PGF, BAT, and liver were obtained after killing by decapitation.

### Serum, liver, and muscle metabolites

Serum insulin levels were determined using the Mouse ultrasensitive insulin ELISA kit (Alpco). Serum FFA levels were measured with Free Fatty acids Quantification kit (BioVision) and serum leptin levels were determined using Mouse/Rat Leptin Quantikine ELISA Kit (R&D systems). Liver and muscle triglyceride content were measured with the Triglyceride Quantification Colorimetric/Fluorometric Kit (BioVision).

### Histology and imaging

SCF, PGF, BAT, and liver were taken and fixed in 4% PFA (Carl Roth, Germany) at room temperature for 24 h dehydrated in an ascending row of ethanol (70%-100%) and xylene. Subsequently, tissues were embedded with paraffin (Leica, Germany) and cut into 2 µm big sections using a microtome (Leica, Germany). Dissected tissue sections were stained with hematoxylin and eosin as previously described (Lillie RD, 1976), and Masson Trichrome staining. Quantification of lipid amount was morphometrically determined by automatic digital image analysis using the commercially available software Definiens Developer XD 2 (Definiens AG, Germany). Immunofluorescent staining was performed on paraffin sliced tissue sections, where antigen retrieval was achieved by boiling with citric acid (pH=6) for 10 min at 800W. After blocking with filtered 3% BSA/PBS at RT for 1h, slides were incubated with primary antibody (Fibronectin 1:50) at 4°C overnight. After subsequent washing (3x PBS, RT, 5min), the slides were incubated with the secondary antibody (anti rabbit 488 (1:200) and DAPI (1:5000)), for 1 hour at RT in the dark. Eventually, the slides were washed (3x PBS, RT, 5min) mounted, cover-slipped and dried for 1h. Imaging was performed with Leica confocal SP5.

### Electron microscopy

Animals were sacrificed and transcardially perfused with PBS followed by a fixative containing 4% paraformaldehyde (Serva, Heidelberg, Germany) and 1.5% glutaraldehyde (Serva). After dissection, the adipose tissue was post-fixed in the same fixative. Next, the fixed tissue was dissected into smaller blocks of approximately 1-2 mm3, which were then further fixed and stained with 0.5% osmium tetroxide (EMS, Hatfield, PA, USA) in PBS for 60 minutes, subsequently rinsed in PBS followed by dehydration with 30%, 50% and 70% ethanol. Afterwards the tissue was stained with 1% uranyl acetate (Merck, Darmstadt, Germany in 70% ethanol for one hour and further dehydrated using 80%, 90%, 96%, and 100% ethanol and finally with propylene oxide (Sigma Aldrich, Steinheim, Germany). The samples were transferred in Durcupan (Sigma Aldrich) and embedded in gelatin capsules followed by the polymerization process at 56°C for 48 hours. After polymerization, the blocks of resin were trimmed and semi-thin sections were prepared using an ultra-microtome (Leica Microsystems, Wetzlar, Germany) to identify regions of interest at the level of light microscopy. After finally trimming the blocks of resin, ultra-thin sections were prepared at a thickness of 55 nm. Sections were transferred on formvar-coated copper grids and stained with lead citrate. The analysis was performed using a Zeiss SIGMA electron microscope equipped with a STEM detector and ATLAS software (Zeiss NTS, Oberkochen, Germany).

For analysis of basal lamina thickness, per animal of each group, 19-21 vessels were analyzed. For each vessel, the basal lamina thickness was measured at 15 different measuring points to calculate the mean value per vessel and mouse. As sectioned vessels regularly exhibit parts with tangential sectioning, thereby artificially broadening the basal lamina, the analysis was confined to the thinnest part of the basal lamina of each vessel, only. Endothelial vesicle density was addressed by counting endothelial vesicles in predefined sections of the vascular circumference to calculate the mean value of endothelial vesicles per µm. Again, parts of the vascular circumference containing endothelial nuclei or accumulations of organelles as well as areas of tangential sectioning were excluded from the analysis.

### Cell culture

For isolation of primary cells, SCF from 8 week-old Kind2^fl/fl^ and Kind2^adipo-cre^ mice was excised and digested in 10 ml of digestion mix (DMEM containing 1% BSA and 0.1% collagenase IV), at 37°C and 1000 rpm shaking for 40 min. For mature adipocyte isolation, digested SCF was filtered through a 250 µm strainer and washed with 5 ml of ice cold PBS. The digested mix was left to stand for 15 min, to allow adipocytes floating to the top. The layer of adipocytes was picked and added to 1 ml of Qiazol for subsequent RNA isolation. For isolation of primary preadipocytes, digested BAT was filtered through a 100 µm filter and washed with 5 ml of ice cold PBS. The suspension was centrifuged at 800g RT for 5min. The supernatant was removed and the pellet was re-suspended in 5ml of culture medium and centrifuged at 800g RT for 5min. The supernatant was discarded and the pellet re-suspended in 3ml of culture medium and eventually plated on a 6-well plate. The cells were immortalized using an ecotropic SV40 large T retrovirus produced in HEK 293T cells. Differentiation of preadipocytes was induced by adding DMEM containing 10% FBS, 1% penicillin and streptomycin, 500µM IBMX/0.5NKOH, 5µM dexamethasone/100%ethanol, 125µM indomethacin/DMSO, 100nM insulin, and 1 nM T3 to each well (Day 0). After two days, the induction medium was replaced by freshly prepared differentiation medium (DMEM containing 10%FBS, 1% penicillin and streptomycin, 100nM insulin, and 1 nM T3). This medium was changed every other day until the cells were fully differentiated (Day 8).

### RNA isolation, reverse transcriptase (RT)-PCR and real-time RT-PCR

In liquid nitrogen ground tissues of BAT (20mg), WAT (80-100mg), and liver (20mg) were homogenized in 1ml of Qiazol (Qiagen) using the TissueLyzer II from (Qiagen). Tissue homogenates were centrifuged (4°C, 2500g, 5 min) and the supernatant was added to 200µl of chloroform, vortexed and centrifugated (4°C, 18400g, 20min). 350µl of the RNA containing clear phase was further added to 70% ethanol, mixed well and pipetted to a spin column of the RNeasy Mini Kit (Qiagen) and RNA isolation was further processed according to the manufacture’s instruction. RNA yield was determined using the Nanodrop (Nanodrop 2000, Thermo Fisher Scientific). For the determination of gene expression, cDNA synthesis (0.5-1 µg total RNA, High Capacity cDNA Reverse Transcription Kit, Applied Biosystems) was performed according to the manufacturers’ instructions. qPCR was performed in a CFX384 Touch (Bio-Rad), using 300 nM forward and reverse primers and iTaq Universal SYBR Green supermix (Bio-Rad). The target gene expression was normalized on TATA box binding protein (TBP) expression. The primer sequences are shown in **Extended Table S1**. Differential expression levels were calculated via the ΔΔct method (Pfaffl, 2001)

### Protein extraction and western blot

Tissues or cells were lysed with RIPA buffer (50 mM Tris pH 7.4, 150 mM NaCl, 1 mM EDTA, 1% Triton X-100), containing 0.1% SDS, 0.01% protease-inhibitor, 0.01% phosphatase-inhibitor cocktail II, and 0.01% phosphatase-inhibitor cocktail III (all from Sigma). Samples were incubated on ice for 10 min, and centrifuged for 10 min at 14.000xg at 4 °C. The supernatant was collected, and protein concentration was determined using the Pierce BCA protein assay kit (Thermo Fischer Scientific). Samples were mixed with 4x sample buffer (Life Technologies) containing 2.5% β-mercaptoethanol (Carl Roth), boiled for 5 min at 95 °C, and loaded on an 7.5-10 % Tris gels for SDS-PAGE with Fisher BioReagentsTM EZ-RunTM Prestained Rec Protein Ladder (ThermoFischer Scientific), as the molecular weight marker and transferred to a PVDF Immobilon-PSQ membrane, 0.45 microns (Merck Millipore). Unspecific binding sites were blocked with 5% non-skimmed milk in TBS-T and incubated in primary antibodies: FAK (Cell Signaling Technology D2R2E; 1:1000), phospho-FAK(Tyr397) (Cell Signaling Technology D20B1; 1:1000), Kindlin-2 (Cell Signaling Technology; 1:1000), integrin beta 1 (Cell Signaling Technology; 1:1000), integrin beta 3 (Cell Signaling Technology; 1:1000), cleaved caspase 3 (Cell Signaling Technology; 1:1000), phosphor-p65-NFKB (Cell Signaling Technology; 1:1000), NFKB (Cell Signaling Technology; 1:1000), insulin receptor (Cell Signaling Technology; 1:1000), phosphor-AKT (Ser473) and phosphor-AKT (Thr308), AKT (Cell Signaling Technology; 1:1000), phosphor-ERK (Cell Signaling Technology; 1:1000), ERK (Cell Signaling Technology; 1:1000), phosphor-p70-S6K (Cell Signaling Technology; 1:1000), p70-S6K (Cell Signaling Technology; 1:1000), UCP1 (Cell Signaling Technology; 1:1000), and α-tubulin (Sigma; 1:4000) at 4 °C over night. After washing, secondary antibodies (anti-rabbit HRP (Cell Signaling Technology; 1:5000), anti-mouse HRP (Cell Signaling Technology; 1:10.000) were incubated for 1 h at RT. The HRP-linked ß-actin (Santa Cruz Biotechnology; 1:5000) was incubated 1 h at RT. After washing, ECL (Merck Millipore) was added to the membranes and the signal was detected using Bio-Rad ChemiDoc™ Imagers (Bio-Rad) or with films (ECL high performance chemiluminescence film (GE Healthcare Life Sciences) or CEA® RP New Medical X-Ray Screen Film Blue Sensitive (CEA Group). Quantifications were performed using Image J software.

### Co-immunoprecipitation assay

Protein was extracted and lysed in NP-40 buffer [1% NP-40, 20 mM Tris-HCl, pH 7.5, 150 mM NaCl, 1 Mm CaCl_2_, 1 mM MgCl_2_ and 10 % glycerol] containing 1 % protease-inhibitor (Sigma-Aldrich) and then centrifuged for 30 min at 14.000 g, 4 °C. Cleared lysate was taken and protein concentration was determined by BCA assay. 4 mg total protein was incubated with 2 µg of rabbit insulin receptor antibody (Cell Signaling Technology, #3025) or 2 µg rabbit IgG control (Cell Signaling Technology, #2729) for 4 hours at 4 °C with rotation. Then, co-immunoprecipitated proteins were purified with µ columns (Miltenyi #130-042-701) according to manufactures protocol. In brief, 50 µl µMACS Protein G MicroBeads (Miltenyi #130-071-101) were added per approach and incubated for 1 h at 4 °C with rotation. All further steps were performed at 4 °C. µcolumns were equilibrated with lysis buffer without glycerol (see above) and lysate was applied on columns. Columns were washed four times with lysis buffer without glycerol and one time with DPBS. Afterwards co-immunoprecipitated proteins were eluted in 50 µl preheated (95 °C) 1X SDS sample buffer (NuPAGE LDS Sample Buffer, #NP0008) supplemented with 2.5% 2-mercaptoethanol and analyzed by western blot. For SDS-gelelectrophoresis 4 to 12 % gels (Thermo Fisher Scientific) were used and electrophoresis was performed at 200 V in MOPS Buffer for 45 min. Blotting was performed in Tris (25 mM), Glycin (192 mM) Buffer at 350 mA for 1.5 h on ice. For blocking, membrane was incubated for 1 hour with 5% non-skimmed milk in TBS-T at RT. Membrane was incubated O/N with rabbit integrin beta 1 antibody (1:1000, Cell Signaling Technology #9699), rabbit integrin beta 3 (1:1000, Cell Signaling Technology #13166) and mouse IR antibody (1:1000, Cell Signaling Technology #3020S) in 5% BSA at 4 °C. On the next day, membrane was washed three times with TBS-Tween20 0.1% for 5 min at RT. After washing, membrane was incubated in secondary antibody either anti-rabbit-IgG-HRP (1:5000, Cell Signaling Technology #7074) or anti-mouse IgG-HRP (1:5000, Santa Cruz Biotechnology #2005) in 5% milk TBS-Tween20 0.1% for 1h at RT. After 3 additional washing steps, each for 5 min washed with TBS-Tween20 0.1%, ECL (Supersignal, West Dura, Thermo Fisher Scientific) was added to the membranes and the signal was detected using Bio-Rad ChemiDoc™ Imagers (Bio-Rad).

### Glucose uptake assay

For glucose uptake, murine white adipose tissue was dissected and digested for 30 minutes at 37°C in Krebs-Ringer buffer (20 mM sodium chloride, 5 mM potassium chloride, 2 mM calcium chloride, 1 mM magnesium chloride, 25 mM sodium bicarbonate and 20 mM HEPES) containing 1 mg/ml collagenase type IV (Gibco) and 1 % BSA (Albumin fraction V, Roth). The digestion was stopped by washing with PBS (Gibco) containing 1 % BSA, and it was in repose for 20 min to let the adipocytes floating. Adipocytes were incubated with or without insulin in the appropriate concentration (1, 10, and 100 nM) for 20 min at 37°C. Subsequently, adipocytes were incubated with 2-deoxy-d-glucose (2-DOG) (Sigma) for 10 min at 37°C (on ice for control) in the presence of the absence of insulin. Glucose uptake was assayed in samples and normalized to protein concentration (Glucose uptake colorimetric kit, Sigma). Results are shown as 2-DG6P uptake in pM per mg protein per min.

### Evans blue staining

To obtain detailed information of the mouse BAT vascular profile, animals were anesthetized with a mixture of ketamine (70 mg/kg) and xylazine (5 mg/kg) transcardially perfused with 0.05 % Evans blue (Sigma E2129) in 5 mL phosphate-buffered saline (PBS) (Gibco, pH 7.4). Perfusions were finalized with 20 mL of 4% paraformaldehyde (PFA) in PBS, pH 7.4, and adipose tissues (SCF, PGF, and BAT) were removed and post-fixed in 4% PFA at 4°C. In either case, adipose tissues were then equilibrated with 30% sucrose in Tris-buffered saline (TBS, pH 7.2) for 48 h before being sectioned into 30 μm coronal slices using a cryostat (CM3050S; Leica, Germany). Eventually, the slides were mounted, cover-slipped and dried for 1h. Imaging was performed with optical microscope.

### Fluorescent angiography

To obtain detailed information of the mouse BAT vascular profile, animals were sacrificed with CO_2_ and transcardially perfused with 20 mL phosphate-buffered saline (PBS) (Gibco, pH 7.4) supplemented with Lectin-FITC conjugate (25 μg/ml; Tritium vulgaris; L4895; Merck, Germany) by using a peristaltic pump at 120 mmHG (Instech, High Flow P720 equipped with 21G canula). Perfusions were finalized with 20 mL of 4% paraformaldehyde (PFA) in PBS, pH 7.4, BAT was removed and post-fixed in 4% PFA at 4°C. In either case, adipose tissues were then equilibrated with 30% sucrose in Tris-buffered saline (TBS, pH 7.2) for 48 h before being sectioned into 30 μm coronal slices using a cryostat (CM3050S; Leica, Germany). Eventually, the slides were mounted, cover-slipped and dried for 1h. Imaging was perfomed with Leica confocal SP5.

### Study in human subjects

We included a cohort of 635 individuals from the Leipzig Biobank (432 women and 203 men) to represent a wide range of BMI (14.7–88.8 kg/m^2^), categories of lean (BMI < 25 kg/m^2^, n=84), overweight (BMI 25.1–29.9 kg/m^2^, n=47), or obesity (BMI > 30 kg/m^2^, n=504). For Pearson’s correlation between both β1 and β3 integrin gene expression in VAT and SAT and body fat percentage (%), 123 individuals were selected for whom complete data sets were available (74 women and 49 men), categories of lean (BMI < 25 kg/m^2^, n=24), overweight (BMI 25.1–29.9 kg/m^2^, n=4), and obese (BMI > 30 kg/m^2^, n=95). From those 123 individuals, 26 individuals had impaired fasting glycemia (n=8), impaired glucose tolerance (n=14) or insulin resistance (n=4). For correlation analyses between HOMA-IR and both β1 and β3 integrin gene expression, 115 individuals without type 2 diabetes with a HOMA-IR range between 0.02 and13.3 were analyzed. Partial correlation, adjusted by body fat (%), between both β1 and β3 integrins gene expression in VAT and SAT and insulin resistance estimated from HOMA-IR was carried out in these individuals (n=115). Collection of human biomaterial, serum analyses and phenotyping were approved by the ethics committee of the University of Leipzig (approval numbers: 159-12-21052012 and 017-12-23012012), and all individuals gave written informed consent before taking part in the study.

### Statistics

All statistics were calculated using Microsoft Excel and GraphPad Prism 8. Data are presented as mean ± standard error of the mean (SEM) unless stated differently in the figure legend. Statistical significance was determined by unpaired Student’s t-test or, for multiple comparisons, using One- or Two-Way ANOVA, followed by Tukey’s multiple comparison’s test, or as stated in the respective figure legend. Differences reached statistical significance with p<0.05. Correlations were established based on Pearson correlation tests for linear relationship with the variants. Correlation coefficient was provided and the *p-*value and *r* value calculated. p<0.05 was adopted as significant.

## Acknowledgements

This work was supported by the project Aging and Metabolic Programming (AMPro). FJRO was supported by a grant to postdoctoral researchers at foreign Universities and Research Centers from the “Alfonso Martín Escudero Foundation”, Spain.

## Author contributions

FJRO, JW and SU designed and conducted the experiments. FJRO and SU wrote the manuscript. MK conducted the electron microcopy experiments. TB, SZ, TG and SR performed experiments. AF conducted size distribution analysis of adipocytes. CGC, TJS, TM, RF and MM designed experiments. MB provided the human data analysis.

## Supporting information

**Expanded View table 1.**
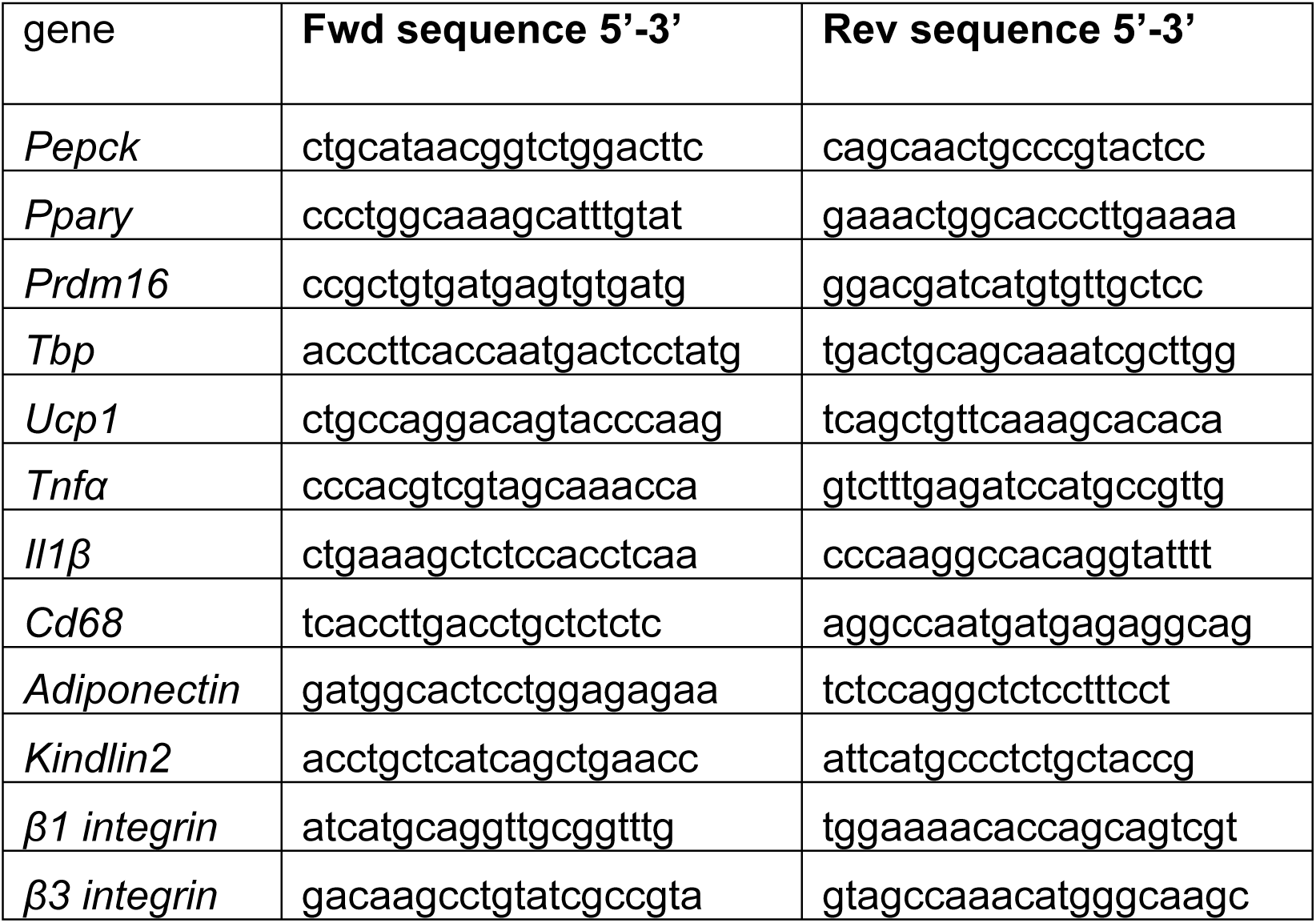
Primer sequences

### Expanded View Figure Legends

**Figure Expanded View 1.**
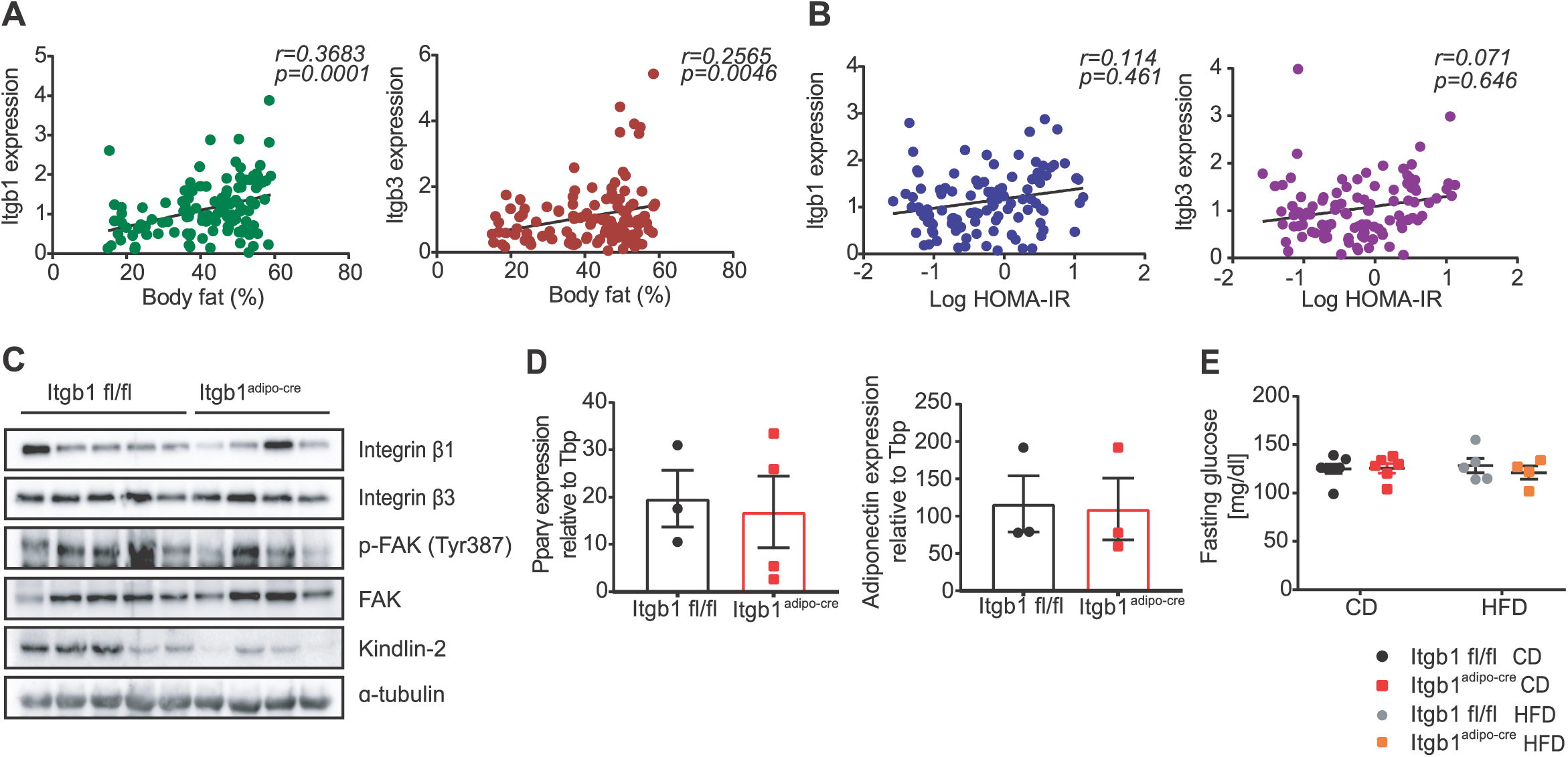
(**A**) Pearson correlation between the both beta 1 (β1) and beta 3 (β3) integrins in subcutaneous adipose tissue (SAT) and body fat (%) in subjects of a range of BMI (n=123). (**B**) Partial correlation, adjusted by body fat (%), between both β1 and β3 integrins in SAT and insulin resistance measured by HOMA-IR in subjects without type 2 diabetes (n=115). (**C**) Representative Western blot images for integrin β1, integrin β3, phospho(Tyr397)- and total FAK, Kindlin-2, and α-tubulin in whole lysate of SCF from integrin beta 1 flox mice (Itgb1 fl/fl) and Itgb1 flox; adiponectin-cre (Itgb1^adipo-cre^) mice fed CD for 13 weeks (n=5-4). (**D**) Relative gene expression of *Pparg* and *adiponectin* in isolated adipocytes of SCF from Itgb1 fl/fl and Itgb1^adipo-cre^ mice fed CD for 8 weeks (n=4). (**E**) Fasting glucose levels (mg/dl) in Itgb1 fl/fl and Itgb1^adipo-cre^ mice fed CD (n=7-6) and HFD for 14 weeks (n=5-4). Data are shown as mean ± SEM. Statistics were calculated using ordinary two-way ANOVA with Tukey’s multiple comparison post-hoc test.

**Figure Expanded View 2.**
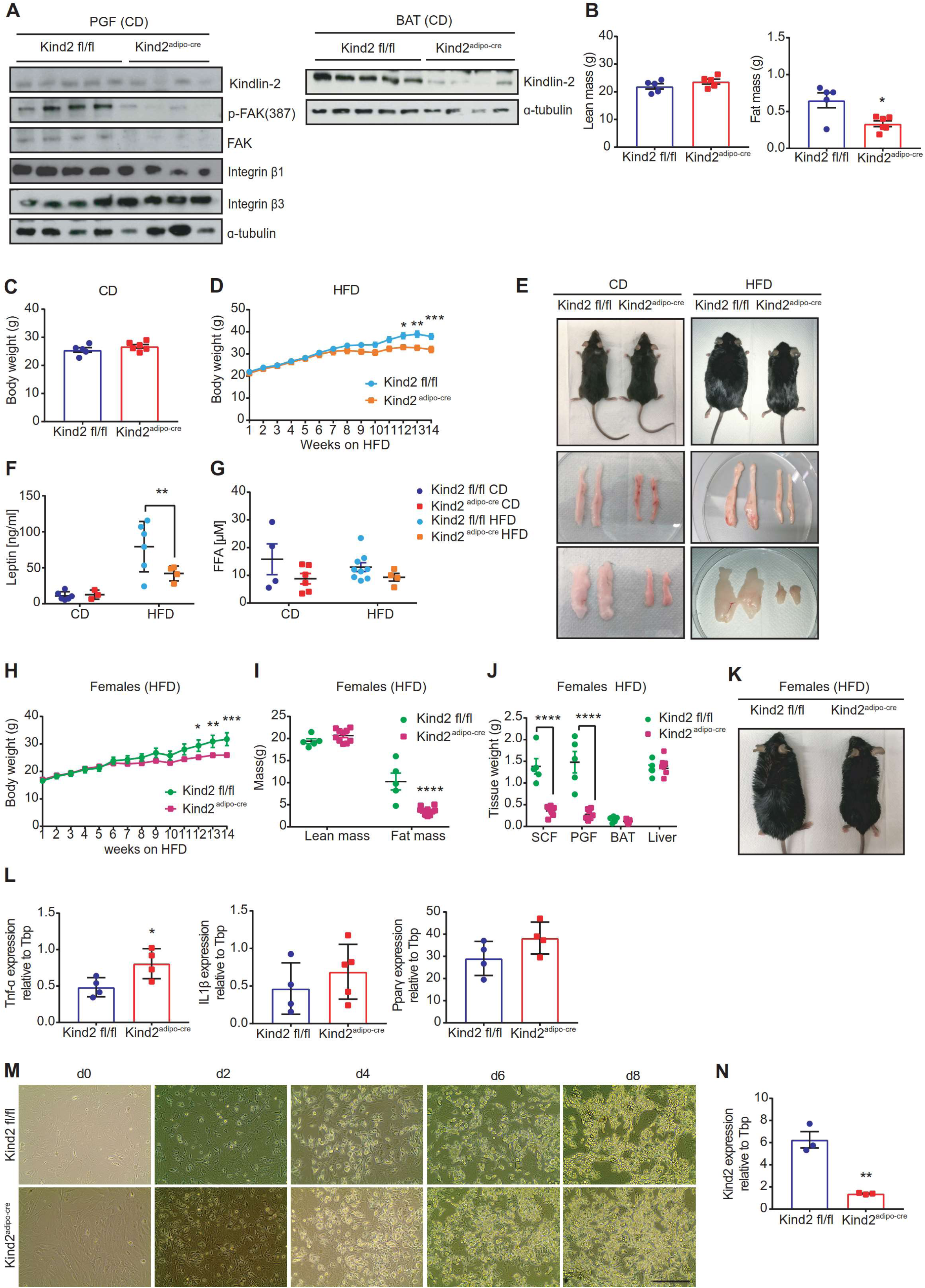
(**A**) Representative western blot images for Kindlin-2, phospho(Tyr397)- and total FAK, integrin β1, integrin β3 and α-tubulin in perigonadal adipose tissue (PGF) and Kindlin-2 and α-tubulin in brown adipose tissue (BAT) from Kindlin-2 flox mice (Kind2 fl/fl) and Kindlin-2 flox; adiponectin-cre (Kind2^adipo-cre^) mice fed CD for 14 weeks. (**B**) Body composition represented as lean and fat mass (g) of Kind2 fl/fl and Kind2^adipo-cre^ mice fed CD at 8 weeks-old (n=5-6). (**C**) Body weight (g) of Kind2 fl/fl and Kind2^adipo-cre^ mice fed CD at 13 weeks-old (n=5-6). (**D**) Body weight development of Kind2 fl/fl and Kind2^adipo-cre^ mice fed HFD for 14 weeks (n=9-4). (**E**) Photograph of Kind2 fl/fl and Kind2^adipo-cre^ mice, and SCF and PGF fed CD and HFD for 14 weeks. (**F**) Serum leptin levels (ng/ml) of Kind2 fl/fl and Kind2^adipo-cre^ mice fed CD (n=5-6) or HFD for 14 weeks (n=9-4). (**G**) Serum free fatty acids (µM) of Kind2 fl/fl and Kind2^adipo-cre^ mice fed CD (n=5-6) or HFD for 14 weeks (n=9-4). (**H**) Body weight development of Kind2 fl/fl and Kind2^adipo-cre^ female mice fed HFD for 14 weeks (n=5-10). (**I**) Body composition of Kind2 fl/fl and Kind2^adipo-cre^ female mice fed HFD for 14 weeks (n=5-10). (**J**) Tissue weights of SCF, PGF, BAT and liver of Kind2 fl/fl and Kind2^adipo-cre^ female mice fed HFD for 14 weeks (n=5-10). (**K**) Photograph of Kind2 fl/fl and Kind2^adipo-cre^ female mice fed HFD for 14 weeks. (**L**) Relative gene expression of *Tnf-α, Il1-β*, and *Pparg* primary adipocytes isolated from SCF from Kind2 fl/fl and Kind2^adipo-cre^ mice fed CD for 8 weeks (n=4). (**M**) Images from primary brown differentiated adipocytes from Kind2 fl/fl and Kind2^adipo-cre^ mice fed CD for 8 weeks (scale bar represents 100 µm). (**N**) Relative gene expression of *Kindlin-2* in primary differentiated brown adipocytes isolated from BAT from Kind2 fl/fl and Kind2^adipo-cre^ mice fed CD for 8 weeks (n=3). Data are shown as mean ± SEM. Statistics were calculated using ordinary two-way ANOVA with Tukey’s multiple comparison post-hoc test (****P<0.0001, *** P<0.001, ** P<0.01, * P<0.05).

**Figure Expanded View 3.**
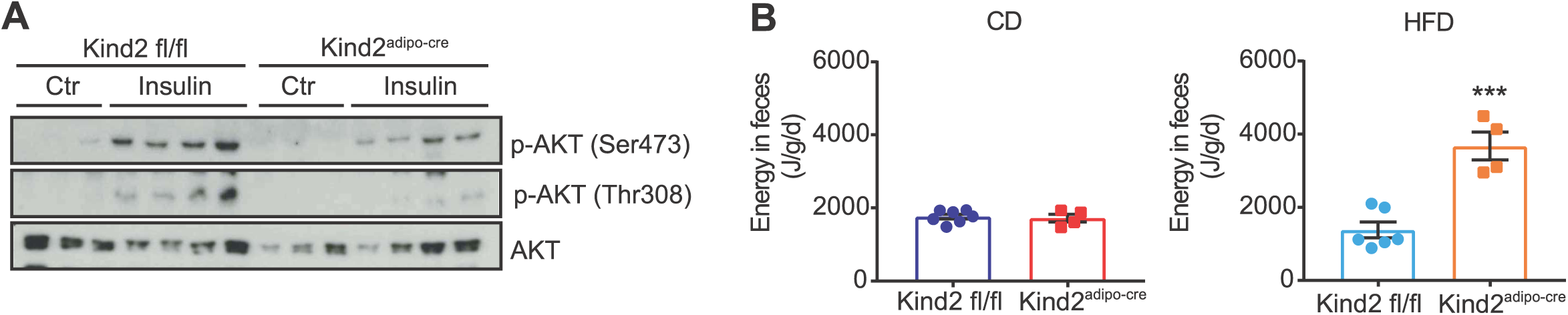
(**A**) Western blot images for phospho-AKT (Ser473), phospho-AKT (Thr308), and AKT in primary adipocytes isolated from perigonadal adipose tissue (PGF) of Kind2 fl/fl and Kind2^adipo-cre^ mice fed CD for 8 weeks in the presence or absence (control=ctr) of insulin [10 nM] for 10 min (n=3). (**B**) Fecal energy content (J/g/d) in Kindlin-2 flox mice (Kind2 fl/fl) and Kindlin-2 flox; adiponectin-cre (Kind2^adipo-cre^) mice fed CD (n=7-4) and HFD (n=6-4) for 14 weeks. Data are shown as mean ± SEM. Statistics were calculated using two-way ANOVA with Tukey’s multiple comparison post-hoc test (***P<0.001, ** P<0.01, * P<0.05).

**Figure Expanded View 4.**
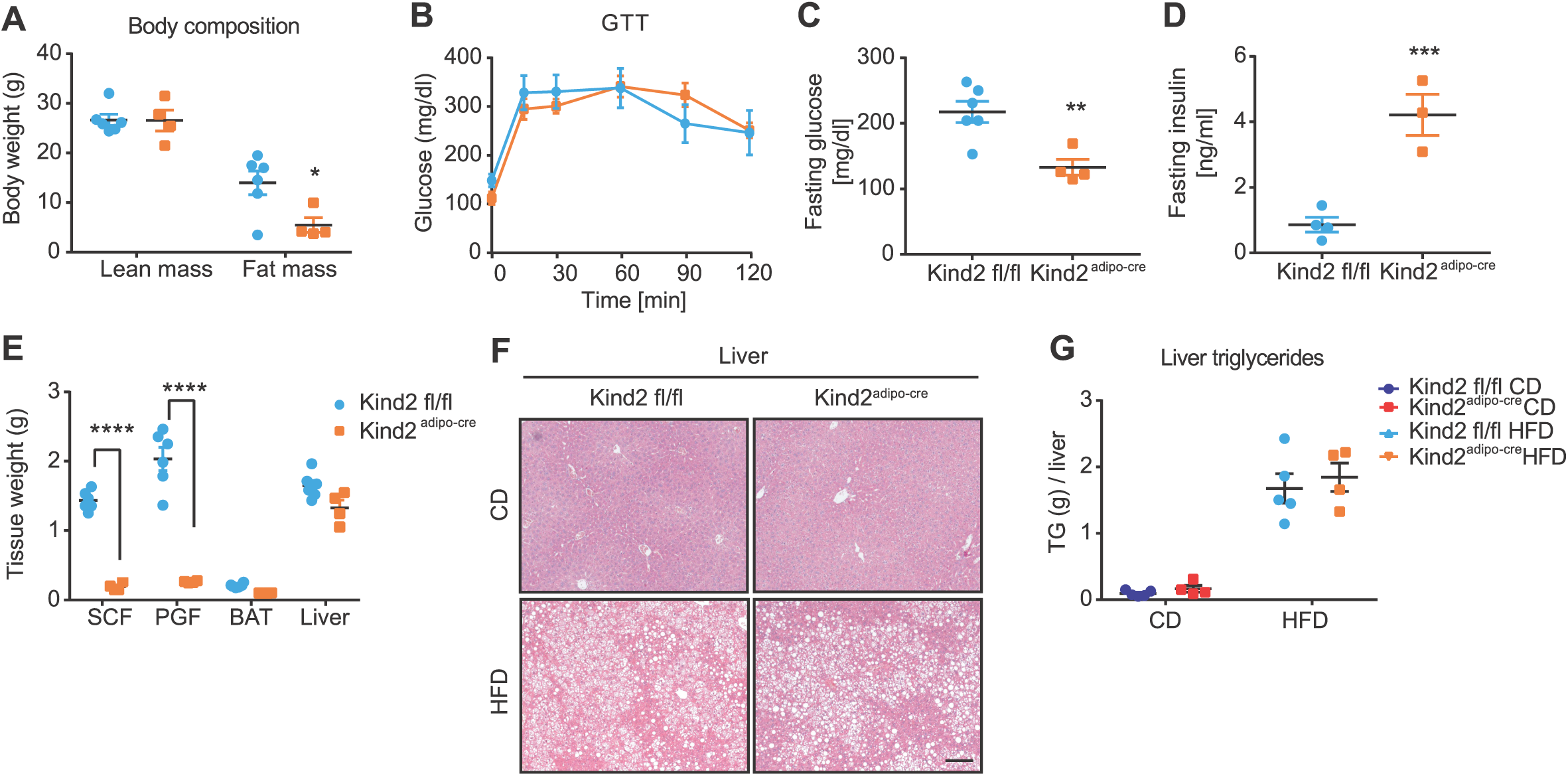
(**A**) Body composition represented as lean and fat mass (g) of Kindlin-2 flox mice (Kind2 fl/fl) and Kindlin-2 flox; adiponectin-cre (Kind2^adipo-cre^) mice fed HFD for 20 weeks (n=6-4). (**B**) Plasma glucose concentration during intraperitoneal glucose tolerance test (2 g/kg) in Kindlin-2 fl/fl and Kind2^adipo-cre^ mice fed HFD for 20 weeks (n=6-4). (**C**) Fasting glucose levels (mg/dl) in Kind2 fl/fl and Kind2^adipo-cre^ mice fed and HFD for 20 weeks (n=6-4). (**D**) Fasting insulin levels (ng/ml) in Kind2 fl/fl and Kind2^adipo-cre^ mice fed HFD for 20 weeks (n=6-4). (**E**) Tissue weights of SCF, PGF, BAT, and liver of Kind2 fl/fl and Kind2^adipo-cre^ female mice fed HFD for 20 weeks (n=6-4). (**F**) Representative H&E staining of liver sections from Kind2 fl/fl and Kind2^adipo-cre^ mice fed CD and HFD for 20 weeks (scale bar represents 100 µm). (**G**) Triglyceride content (g) per whole liver in Kindlin-2 fl/fl and Kind2^adipo-cre^ mice fed CD (n=4) and HFD for 20 weeks (n=5-4). Data are shown as mean ± SEM. Statistics were calculated using two-way ANOVA with Tukey’s multiple comparison post-hoc test (****P<0.0001, ***P<0.001, ** P<0.01, * P<0.05).

**Figure Expanded View 5.**
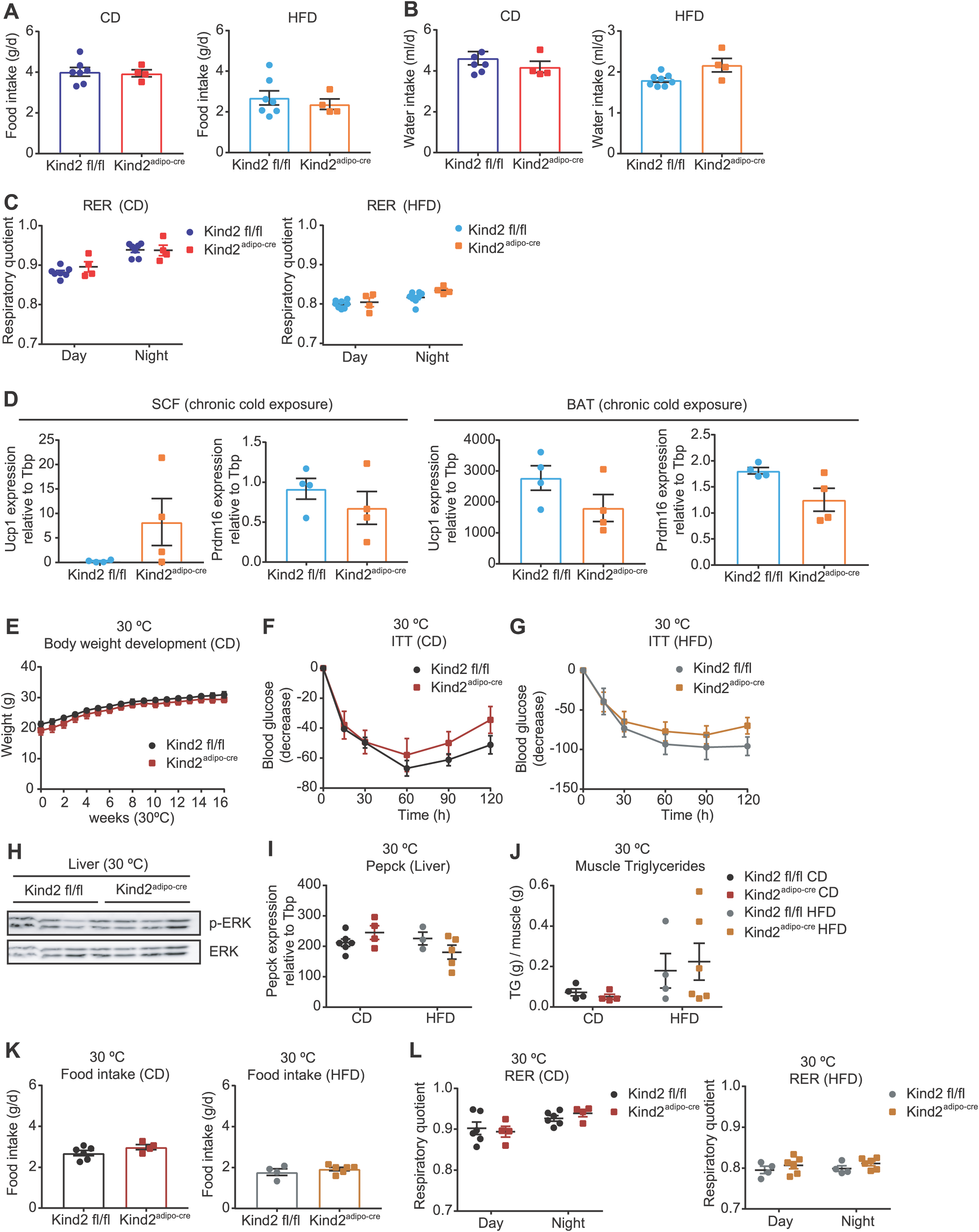
(**A**) Food intake (g) per day in Kindlin-2 flox (Kind2 fl/fl) and Kindlin-2 flox; adiponectin-cre (Kind2^adipo-cre^) mice fed CD (n=7-4) and HFD (n=6-4) for 14 weeks. (**B**) Water intake (ml) per day in Kind2 fl/fl and Kind2^adipo-cre^ mice fed CD (n=6-4) and HFD (n=6-4) for 14 weeks. (**C**) Respiratory quotient (RER) average for 3 days in Kind2 fl/fl and Kind2^adipo-cre^ mice fed CD (n=7-4) and HFD (n=6-4) for 14 weeks. (**D**) Relative gene expression of *Ucp1* and *Prdm16* in subcutaneous adipose tissue (SCF) and brown adipose tissue (BAT) from Kind2 fl/fl and Kind2^adipo-cre^ mice fed HFD for 14 weeks after chronic cold exposure at 4 °C for 48 h (n=4). (**E**) Body weight development of Kind2 fl/fl and Kind2^adipo-cre^ mice fed CD for 16 weeks housed at thermoneutrality (30°C) (n=6-4). (**F**) Insulin tolerance test (ITT) after i.p. injection of 0.75 IU/kg of insulin in Kind2 fl/fl and Kind2^adipo-cre^ mice fed CD (n=6-4) for 16 weeks housed at thermoneutrality (30°C). (**G**) ITT after i.p. injection of 1.25 IU/kg in Kind2 fl/fl and Kind2^adipo-cre^ mice fed HFD (n=4-6) for 16 weeks housed at thermoneutrality (30°C). (**H**) Representative western blot images for phospho-ERK and ERK in liver from Kind2 fl/fl and Kind2^adipo-cre^ mice fed HFD for 16 weeks housed at thermoneutrality (30°C) (n=4-3). (**I**) Relative gene expression of *Pepck* in liver from Kind2 fl/fl and Kind2^adipo-cre^ mice fed CD (n=6-4) and HFD (n=3-5) for 16 weeks housed at thermoneutrality (30°C). (**J**) Triglyceride content in muscle (g/g) in Kindlin-2 fl/fl and Kind2^adipo-cre^ mice fed CD (n=4) and HFD for 16 weeks housed at thermoneutrality (30°C) (n=3-6). (**K**) Food intake (g) per day in Kind2 fl/fl and Kind2^adipo-cre^ mice fed CD (n=6-4) and HFD (n=4-6) for 16 weeks housed at thermoneutrality (30°C). (**L**) Respiratory quotient (RER) average for 3 days separated by day and night cycles in Kind2 fl/fl and Kind2^adipo-cre^ mice fed CD (n=6-4) and HFD (n=4-6) for 16 weeks housed at thermoneutrality (30°C). Data are shown as mean ± SEM. Statistics were calculated using two-way ANOVA with Tukey’s multiple comparison post-hoc test (***P<0.001, ** P<0.01, * P<0.05).

**Figure Expanded View 6.**
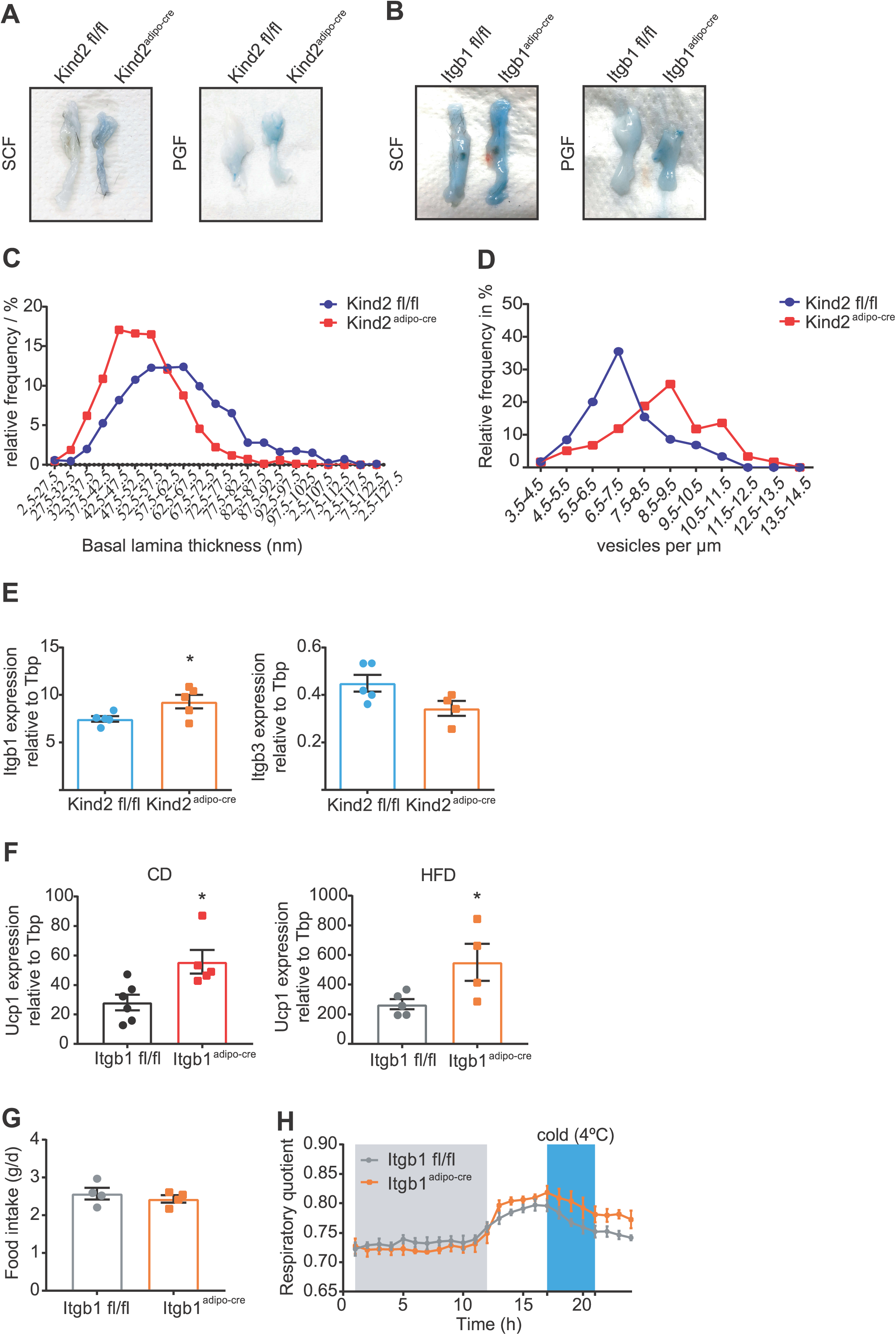
(**A**) Photograph of subcutaneous (SCF) and perigonadal (PGF) adipose tissue from Kindlin-2 flox (Kind2 fl/fl) and Kindlin-2 flox; adiponectin-cre (Kind2^adipo-cre^) mice fed CD for 8 weeks after Evans blue (0.05 % in PBS) perfusion and fixation with 4 % PFA. (**B**) Photograph of SCF and PGF adipose tissue from integrin β1 flox mice (Itgb1 fl/fl) and Itgb1 flox; adiponectin-cre (Itgb1^adipo-cre^) mice fed CD for 8 weeks after Evans blue (0.05 % in PBS) perfusion and fixation with 4 % PFA. (**C**) Values distribution of basal lamina thickness quantification (µm) measured by EM at 15 different points to calculate the mean value per vessel and mouse in BAT from Kind2 fl/fl and Kind2^adipo-cre^ mice fed CD for 8 weeks (n=3). (**D**) Frequency distribution of endothelial vesicle density (per µm) measured by EM in BAT from Kind2 fl/fl and Kind2^adipo-cre^ mice fed CD for 8 weeks (n=3). (**E**) Relative gene expression of *Itgb1* and *Itgb3* in brown adipose tissue (BAT) from Kind2 fl/fl and Kind2^adipo-cre^ mice fed HFD for 14 weeks (n=5-4). (**F**) Relative gene expression of *Ucp1* in BAT from Itgb1 fl/fl and Itgb1^adipo-cre^ mice fed CD (n=6-5) and HFD for 14 weeks (n=5-4). (**G**) Food intake (g) per day in Itgb1 fl/fl and Itgb1^adipo-cre^ mice fed HFD for 14 weeks (n=5-4). (**H**) Respiratory quotient (RER) average of 3 days in Itgb1 fl/fl and Itgb1^adipo-cre^ mice fed HFD (n=5-4) for 14 weeks during acute cold exposure at 4 °C for 5 hours (n=5-4). Data are shown as mean ± SEM. Statistics were calculated using two-way ANOVA with Tukey’s multiple comparison post-hoc test (** P<0.01, * P<0.05).

## References

Boucher J KA, Kahn CR. (2014) Insulin receptor signaling in normal and insulin-resistant states. Cold Spring Harb Perspect Biol 6

Bui T, Rennhack J, Mok S, Ling C, Perez M, Roccamo J, Andrechek ER, Moraes C, Muller WJ (2019) Functional Redundancy between beta1 and beta3 Integrin in Activating the IR/Akt/mTORC1 Signaling Axis to Promote ErbB2-Driven Breast Cancer. Cell Rep 29: 589–602 e6

Calderwood DAC, I. D.; Critchley, D. R. (2013) Talins and kindlins: partners in integrin-mediated adhesion. Nat Rev Mol Cell Biol 14: 503–17

Cannon B NJ (2004) Brown adipose tissue: function and physiological significance. Physiol Rev 84: 277–359

Charo IF NL, Smith JW, Cheresh DA (1990) The vitronectin receptor alpha v beta 3 binds fibronectin and acts in concert with alpha 5 beta 1 in promoting cellular attachment and spreading on fibronectin. J Cell Biol 111

Chloé C. Féral JGN, Christiane Kummer, Marina Slepak, Jerrold M. Olefsky, Mark H. Ginsberg (2008) Blockade of alpha4 integrin signaling ameliorates the metabolic consequences of high-fat diet-induced obesity. Diabetes 57: 1842–51

Clayton ZS MC (2018) Short-term thermoneutral housing alters glucose metabolism and markers of adipose tissue browning in response to a high-fat diet in lean mice. American Journal of Physiology- Regulatory, Integrative and Comparative Physiology 315: R627–R637

Cui X NN, Zarebidaki E, Cao Q, Li F, Zha L, Bartness T, Shi H, Xue B (2016) Thermoneutrality decreases thermogenic program and promotes adiposity in high-fat diet-fed mice. Physiological Reports 4

Dibble CC, Manning BD (2013) Signal integration by mTORC1 coordinates nutrient input with biosynthetic output. Nat Cell Biol 15: 555–64

Gao H, Guo Y, Yan Q, Yang W, Li R, Lin S, Bai X, Liu C, Chen D, Cao H, Xiao G (2019) Lipoatrophy and metabolic disturbance in mice with adipose-specific deletion of kindlin-2. JCI Insight 4

Guilherme A TK, Czech MP (1998) Cross-talk between insulin receptor and integrin alpha5 beta1 signaling pathways. J Biol Chem 273

Harburger DS, Bouaouina M, Calderwood DA (2009) Kindlin-1 and −2 directly bind the C-terminal region of beta integrin cytoplasmic tails and exert integrin-specific activation effects. J Biol Chem 284: 11485–97

Heemskerk JW, Mattheij NJ, Cosemans JM (2013) Platelet-based coagulation: different populations, different functions. J Thromb Haemost 11: 2–16

Horton M (1997) The αvβ3 integrin “vitronectin receptor”. Int J Biochem Cell Biol 29

Hynes R (2002) Integrins: bidirectional, allosteric signaling machines. Cell 110: 673–87

Kahn CR WG, Lee KY (2019) Altered adipose tissue and adipocyte function in the pathogenesis of metabolic syndrome. J Clin Invest 129: 3990–4000

Kang L AJ, Lee-Young RS, Zhang Z, James FD, Neufer PD, Pozzi A, Zutter MM, Wasserman DH. (2011) Diet-induced muscle insulin resistance is associated with extracellular matrix remodeling and interaction with integrin alpha2beta1 in mice. Diabetes 60: 416–26

Kloeker S, Major MB, Calderwood DA, Ginsberg MH, Jones DA, Beckerle MC (2004) The Kindler syndrome protein is regulated by transforming growth factor-beta and involved in integrin-mediated adhesion. J Biol Chem 279: 6824–33

Lee PL, Tang Y, Li H, Guertin DA (2016) Raptor/mTORC1 loss in adipocytes causes progressive lipodystrophy and fatty liver disease. Mol Metab 5: 422–432

Lillie RD PP, Donaldson PT. (1976) Hematoxylin substitutes: a survey of mordant dyes tested and consideration of the relation of their structure to performance as nuclear stains. Stain Technol 51: 25–41

Luk CT, Shi SY, Cai EP, Sivasubramaniyam T, Krishnamurthy M, Brunt JJ, Schroer SA, Winer DA, Woo M (2017) FAK signalling controls insulin sensitivity through regulation of adipocyte survival. Nat Commun 8: 14360

Luo L, Liu, M. (2016) Adipose tissue in control of metabolism. J Endocrinol 231: R77–R99

Ma YQ, Qin J, Wu C, Plow EF (2008) Kindlin-2 (Mig-2): a co-activator of beta3 integrins. J Cell Biol 181: 439–46

Montanez E, Ussar S, Schifferer M, Bosl M, Zent R, Moser M, Fassler R (2008) Kindlin-2 controls bidirectional signaling of integrins. Genes Dev 22: 1325–30

Ojakian GK SR (1994) Regulation of epithelial cell surface polarity reversal by β1 integrins. J Cell Sci 107

Peirce V. PV, Vidal-Puig A. (2016) Adipose Structure (White, Brown, Beige). In Metabolic Syndrome, pp 369–396.

Pfaffl M (2001) A new mathematical model for relative quantification in real-time RT-PCR. Nucleic Acids Res 29

Polak P, Cybulski N, Feige JN, Auwerx J, Ruegg MA, Hall MN (2008) Adipose-specific knockout of raptor results in lean mice with enhanced mitochondrial respiration. Cell Metab 8: 399–410

Potocnik AJ BC, Fässler R. (2000) Fetal and adult hematopoietic stem cells require beta1 integrin function for colonizing fetal liver, spleen, and bone marrow. Immunity 12: 653–63

Qiang GWK, H.; Xu, S.; Pham, H. A.; Parlee, S. D.; Burr, A. A.; Gil, V.; Pang, J.; Hughes, A.; Gu, X.; Fantuzzi, G.; MacDougald, O. A.; Liew, C. W. (2016) Lipodystrophy and severe metabolic dysfunction in mice with adipose tissue-specific insulin receptor ablation. Mol Metab 5: 480–490

Schneller M VK, Ruoslahti E. (1997) Alphavbeta3 integrin associates with activated insulin and PDGFbeta receptors and potentiates the biological activity of PDGF. EMBO JOURNAL 16: 5600–7

Schoettl T, Fischer IP, Ussar S (2018) Heterogeneity of adipose tissue in development and metabolic function. J Exp Biol 221

Shi X, Ma YQ, Tu Y, Chen K, Wu S, Fukuda K, Qin J, Plow EF, Wu C (2007) The MIG-2/integrin interaction strengthens cell-matrix adhesion and modulates cell motility. J Biol Chem 282: 20455–66

Shimobayashi M, Albert V, Woelnerhanssen B, Frei IC, Weissenberger D, Meyer-Gerspach AC, Clement N, Moes S, Colombi M, Meier JA, Swierczynska MM, Jeno P, Beglinger C, Peterli R, Hall MN (2018) Insulin resistance causes inflammation in adipose tissue. J Clin Invest 128: 1538–1550

Sun K, Kusminski CM, Scherer PE (2011) Adipose tissue remodeling and obesity. J Clin Invest 121: 2094–101

Sun K, Tordjman J, Clement K, Scherer PE (2013) Fibrosis and adipose tissue dysfunction. Cell Metab 18: 470–7

Theodosiou M WM, Böttcher RT, Rognoni E, Veelders M, Bharadwaj M, Lambacher A, Austen K, Müller DJ, Zent R, Fässler R (2016) Kindlin-2 cooperates with talin to activate integrins and induces cell spreading by directly binding paxillin. Elife 5: e10130

Ussar S, Wang, H. V., Linder, S., Fassler, R., Moser, M. (2006) The Kindlins: subcellular localization and expression during murine development. Exp Cell Res 312: 3142–51

Vrieze A SJ, Admiraal WM, Soeters MR, Nieuwdorp M, Verberne HJ, Holleman F (2012) Fasting and postprandial activity of brown adipose tissue in healthy men. J Nucl Med 53: 1407–10

Vuori K RE (1994) Association of insulin receptor substrate-1 with integrins. Science 266: 1576–8

Wang HV, Chang, L. W., Brixius, K., Wickstrom, S. A., Montanez, E., Thievessen, I., Schwander, M., Muller, U., Bloch, W., Mayer, U., Fassler, R. (2008) Integrin-linked kinase stabilizes myotendinous junctions and protects muscle from stress-induced damage. J Cell Biol 180: 1037–49

Williams AS, Kang L, Wasserman DH (2015) The extracellular matrix and insulin resistance. Trends Endocrinol Metab 26: 357–66

Wu J, Bostrom P, Sparks LM, Ye L, Choi JH, Giang AH, Khandekar M, Virtanen KA, Nuutila P, Schaart G, Huang K, Tu H, van Marken Lichtenbelt WD, Hoeks J, Enerback S, Schrauwen P, Spiegelman BM (2012) Beige adipocytes are a distinct type of thermogenic fat cell in mouse and human. Cell 150: 366–76

Yoshikazu Takada XY, Scott Simon (2007) The integrins. Genome Biol 8: 215

Yudasaka M, Yomogida Y, Zhang M, Nakahara M, Kobayashi N, Tanaka T, Okamatsu-Ogura Y, Saeki K, Kataura H (2018) Fasting-dependent Vascular Permeability Enhancement in Brown Adipose Tissues Evidenced by Using Carbon Nanotubes as Fluorescent Probes. Sci Rep 8: 14446

